# Repeated losses and gains of bacterial symbionts in gutless marine annelids over 150 million years

**DOI:** 10.1101/2021.04.28.441735

**Authors:** Anna Mankowski, Manuel Kleiner, Christer Erséus, Nikolaus Leisch, Yui Sato, Jean-Marie Volland, Bruno Huettel, Cecilia Wentrup, Tanja Woyke, Juliane Wippler, Nicole Dubilier, Harald R. Gruber-Vodicka

## Abstract

Multipartite symbioses, which involve intimate interactions among three or more species, have evolved independently numerous times in convergent evolution, and enable functional complexity and ecological flexibility. How stability is maintained in such relationships remains, however, poorly understood, and can only be fully understood by integrating processes across both micro- and macroevolutionary scales. In this study, we bridge these scales by examining the symbiotic communities of gutless marine oligochaetes - annelid worms that have become obligately dependent on their bacterial partners for nutrition and waste removal over the past 150 million years. Using metagenomic sequencing of 231 individuals from 63 host species, collected across 17 marine environments worldwide, we reconstructed host and symbiont phylogenies to investigate the composition, specificity, and evolutionary history of their symbiotic communities from the strain to genus level. Our results show that each gutless oligochaete harbors up to 10 symbiont species drawn from a restricted pool of 33 marine bacterial genera. Community composition was strongly shaped by host species at 90% explanatory power, and much less by geography or environment. Symbiont communities were highly stable within host species but shifted rapidly at short macroevolutionary scales as host species diverged. Ancestral state reconstructions revealed that the primary symbiont, *Candidatus* Thiosymbion, was an early and persistent acquisition, while secondary symbionts were gained multiple times through convergent evolution, enhancing the diversity and metabolic flexibility of the symbiotic communities. By linking fine-scale microevolutionary dynamics with broader macroevolutionary patterns, our study reveals how gutless oligochaetes achieve both stability and flexibility in their symbiotic partnerships - enabling long-term host dependency while driving innovation and adaptation. This evolutionary trajectory may underpin the ecological success and diversification of gutless oligochaetes in marine sediments, and may help explain the repeated emergence and persistence of other multipartite associations.

## Main

Most animals are associated with microorganisms, which modulate central features of their host’s biology, ecology and evolution, such as its nutrition^1–14^, development^15–18^, and defense against predators or pathogens^19–21^. In many associations, the microorganisms provide fitness benefits to their hosts, and in some highly intimate symbioses, the host has become obligately dependent on its microbiota for its survival, for example in nutritional symbioses^2,5,22,23^. In these obligate symbioses, high degrees of partner fidelity are often essential for the long-term stability of the association^24–26^, particularly in hosts that rely on a single symbiont species that is strictly vertically transmitted, such as some insects^27,28^ and marine invertebrates^2,29^. However, there are many animals that are intimately and obligately associated with multiple symbiont species, like mealybugs, leafhoppers and deep-sea mussels ^30–34^. In these multipartite and obligate associations, it is not well understood how stable the symbiont communities are over evolutionary time, and which processes are involved in shaping stability.

Studies on symbiont community composition and partner fidelity in animals with multiple microbial partners have commonly focused on either microevolutionary scales, that is within a host species^35–39^ or macroevolutionary scales, that is across host species^4,40–53^. However, macroevolutionary patterns are often shaped by microevolutionary processes, making it essential to connect the two for a comprehensive understanding of how lineages have evolved. Linking these processes in symbiotic associations and disentangling the effects of evolution, ecology, and biogeography requires examining a broad diversity of hosts and symbionts across a wide range of environments and geographic regions. Only a handful of studies have investigated macroevolutionary processes across an extensive phylogenetic and geographic range of a host clade, but to our knowledge, these have not included microevolutionary analyses^40,51,53,54^.

Gutless marine oligochaetes are an ideal group for investigating how stability is maintained in obligate associations with multiple symbionts, as they harbor a diverse, yet tractable number of five to seven co-occurring bacterial species^55–61^. The hosts are phylogenetically diverse with over 100 described species that belong to two genera, *Olavius* and *Inanidrilus*, which have descended from a single common ancestor, forming a monophyletic host clade^62^. Gutless oligochaetes occur around the world in a wide range of environments in mainly tropical to temperate coastal habitats, although some species have also been found in the deep sea ^37,55–61,63^. Molecular analyses indicated that these small, sediment-dwelling annelids of a few mm length and about 0.1-0.5 mm width diverged from their gut-bearing oligochaete relatives more than 50 million years ago (Ma)^64^. Over time, these hosts have become so obligately dependent on their bacterial symbionts for their nutrition and recycling of their waste compounds that they no longer have digestive and excretory systems^55,63,65,66^.

The dominant symbiont in terms of abundance and biomass in the 22 gutless oligochaete species investigated so far is ‘*Candidatus* Thiosymbion’, a chemoautotrophic sulfur oxidizer (from here on *Thiosymbion*)^67^. The less abundant secondary symbionts have been analyzed in only seven gutless oligochaete species: In these host species, four to six bacterial species co-occur consistently in individuals from the same host species, but across host species, symbiont communities are highly variable ^37,55–61,63^. Imaging analyses of symbiont transmission in three gutless oligochaete species revealed that symbionts are passed vertically from one generation to the next through smearing when the eggs are deposited in the sediment^68–71^. However, these studies could not fully resolve if all symbionts are acquired vertically, or if some are acquired horizontally from the environment during egg deposition. In the only gutless oligochaete species in which partner fidelity was investigated in detail, *Olavius algarvensis*, fidelity varied across the full range, from strict in *Thiosymbion* to moderate or absent across the different secondary symbiont species, indicating both vertical and horizontal modes of transmission depending on the symbiont lineage^37^.

Decades of extensive sampling of gutless oligochaetes around the world have now provided a dataset that has enough phylogenetic, geographic, and environmental diversity to gain insights into the ecological and evolutionary processes that underpin these symbioses. Using Illumina shotgun metagenomic sequencing, we analyzed 231 host individuals from 63 species, collected at 17 sites in the Atlantic, Caribbean, Mediterranean, Red Sea, Indian Ocean and Pacific Ocean. Phylogenetic marker genes for the symbionts and hosts were extracted from the metagenomes, and used to reconstruct their diversity and phylogeny (16S rRNA for the symbionts, nuclear 28S rRNA and mitochondrial cytochrome c oxidase subunit I (mtCOI) for the hosts), test for phylosymbiosis, and reveal how host species, geography, and environment have shaped the symbiotic communities of gutless oligochaetes. We next employed a structured approach spanning microevolutionary to macroevolutionary scales by analyzing specificity and fidelity within and across host species, as well as across symbiont genera, using genome-scale resolution provided by metagenome-assembled genomes (MAGs) of symbionts. Finally, we reconstructed the acquisition and loss of symbiont clades throughout the diversification of gutless oligochaetes. These analyses revealed how community stability at microevolutionary scales, coupled with flexibility over macroevolutionary timescales, has shaped the symbioses in a species-rich group of marine invertebrates from across the globe.

## Results and discussion

### Symbiont communities are recruited from a limited number of marine genera

We reconstructed full-length 16S rRNA genes of the bacterial symbionts from the metagenomes of each of the 231 host individuals from 63 host species, sampled at 17 geographic locations (Fig. S1, Table S1). We first defined symbiont clades at the genus level (> 95% 16S rRNA identity) and focused on genera that were present in at least 50% of the individuals of a host species (Fig. 1, Table S2, Note S1). Based on these criteria, we identified 33 symbiont genera that belonged to six classes – with *Gammaproteobacteria* the most common across host species, followed by *Alpha*- and *Desulfobacterota* (*Deltaproteobacteria*), as well as *Spirochaetia*, *Acidimicrobiia*, and an undefined class of *Marinisomatot*a (*Marinimicrobia*) (Table S2). Fourteen of the 33 symbiont genera were not previously known to be associated with gutless oligochaetes^55,57–61,72^.

**Figure 1:**
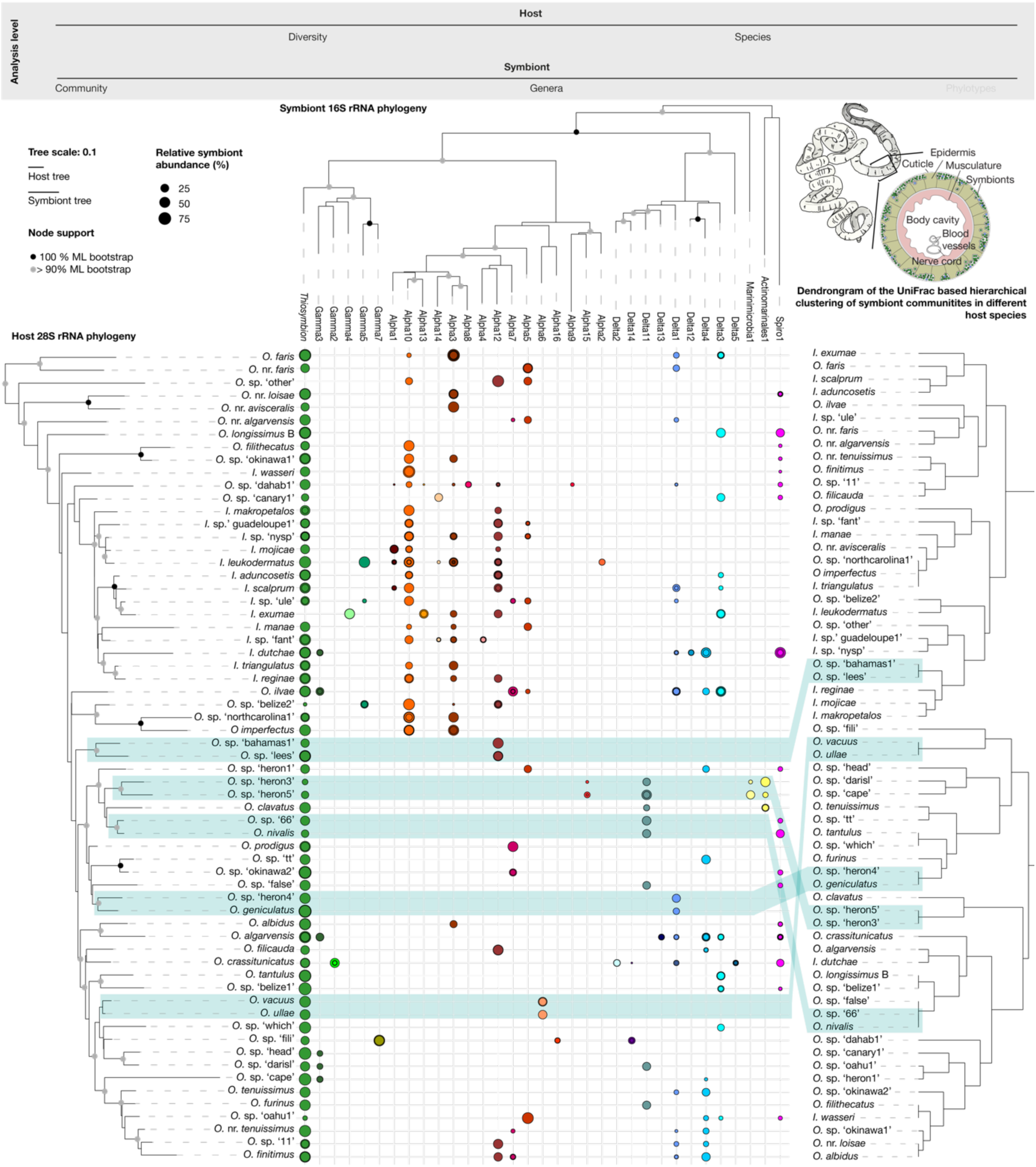
Symbiont communities across 64 gutless oligochaete species are recruited from 33 bacterial genera. **The composition of the symbiotic communities is not shaped by host phylogeny at longer evolutionary timescales, as only some closely related host species share similar symbiont communities** (highlighted in green-blue shading). Top right: Habitus of a gutless oligochaete and a cross section of the worm showing the symbiont-containing region between the cuticle and the epidermis. Trees: Maximum-likelihood phylogeny of the host 28S rRNA gene (left) and symbiont 16S rRNA gene (top). Both host and symbiont trees were calculated with all available sequences. For visualization, one representative sequence per host species/symbiont genus is shown. Scale bar indicates 10% estimated sequence divergence. Bootstrap support > 90% is highlighted in both trees, with the 100%-supported nodes shown in black. Middle panel: Relative abundance of symbiont genera per host species as estimated with EMIRGE. If multiple host individuals of the same species were analyzed, the dots showing their abundance are overlaid. Right tree: Dendrogram of symbiont community composition showing UniFrac distances between host species based on average abundances per host species (Table S4). Congruences between symbiont community composition and host 28S rRNA gene tree are shaded in green-blue.

We next characterized the composition of the symbiont community in each individual and found that each host species was associated with two to ten of the 33 symbiont genera (mean: 3.40 genera, median: 3.00 genera, s.d.: 1.23). For the symbiont genera, the number of hosts species with which they were associated was highly variable. Only one genus, *Thiosymbion*, was present in nearly all host species. Most symbiont genera were present in at least two and up to 21 species, while only a few were found in only a single host species (Fig. 1 and Table S3).

We then examined if gutless oligochaete symbiont genera are associated with other host groups or have been found in environmental samples (e.g., sediments, water, etc.), by analyzing the phylogenetic relationships between all gutless oligochaete symbionts and their closest relatives from publicly available 16S rRNA gene sequences. In addition, we sequenced the metagenomes of the closest relatives of gutless oligochaetes, which have guts and often co-occur with gutless oligochaetes in the same habitats, and screened them for the presence of shared symbiont genera (Fig. S2-S34, S35, Note S2). These analyses revealed that 21 symbiont genera were only associated with gutless oligochaetes. Of the remaining 12 genera, six were also found in association with other hosts that all regularly co-occur with gutless oligochaetes^73^. As previously reported, the genus *Thiosymbion* is also associated with the marine nematodes Stilbonematinae and *Astomonema*^67^. In addition, close relatives of the Gamma5 and Gamma7 symbionts have been found on stilbonematine nematodes (*Eubostrichus topiarius*), the Gamma4 symbionts are members of ‘*Candidatus* Kentron’, the primary symbionts of the marine ciliate *Kentrophoros*^74^, and we found the Alpha3 and Alpha8 symbionts in the gut-bearing relatives of the gutless oligochaetes (Fig. S2-S34, S35). Only six of the 33 symbiont genera were phylogenetically intermixed with bacteria from environmental sources (Alpha14, Delta1, Delta3, Delta4, Delta11 and Gamma3, Fig. S2-S34, Note S3). The predominance of symbionts from host-associated genera in gutless oligochaetes suggests that these bacteria might have acquired traits in their interactions with other hosts that enabled them to easily colonize new hosts. These findings further indicate that symbiont transfers across distantly related marine invertebrate phyla may be more widespread than previously recognized, pointing to a far more interconnected host–microbe network than currently appreciated.

Although we analyzed the symbiont communities of nine times more host species than previously described, we discovered only roughly two times more symbiont genera (63 host species compared to seven, 33 symbiont genera identified compared to 17)^55–61^. This corresponded with our rarefaction analysis showing a saturation in the detection of new symbiont genera, and indicates that the symbionts of gutless oligochaetes are acquired from a limited pool of bacterial groups (Fig. S36). Given the high diversity of microorganisms in marine sediments, the environment in which gutless oligochaetes live^75^, our results suggest that only a very small subset of sediment bacteria are able to associate with marine oligochaetes^52,76–79^.

### The symbiont communities of gutless oligochaetes are largely specific to host species

To understand the processes that shape symbiont community composition, we first screened for symbiont-symbiont interactions that would select either for or against any given combination of symbiont genera, using a co-occurrence network analysis. We did not observe any significant symbiont-symbiont exclusion patterns and found only two significant co-occurrence patterns between two pairs of genera (Delta11 and Actinomarinales1, Alpha 15 and Marinimicrobia1, Fig. S37, Note S4). We then compared the influence of host species, geography, and the environmental factors organic input and sediment type, on symbiont community composition (Fig. 2, Table S3). Host species had by far the highest explanatory power for differences in symbiont community composition at 90%, compared to 20% for ocean basin, and 7 and 13% for organic input and sediment type respectively (analysis based on UniFrac distances between symbiont communities; PERMANOVA: host species: 90.05%, p=0.001, ocean basin: 19.85%, p=0.001, organic input: 6.59%, p=0.011, sediment type: 13.02%, p=0.001, Mantel test for geographic distance: R=0.18, p=0.001, Fig. 2).

**Figure 2:**
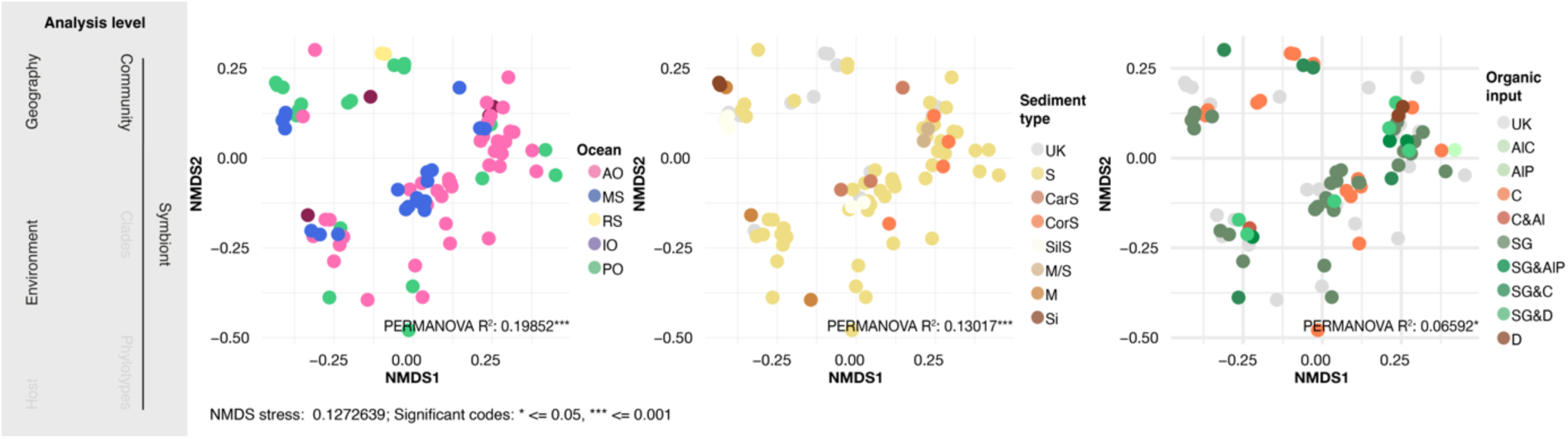
Symbiont community composition is not as strongly shaped by ocean basin (20%) and environmental parameters (7 and 13% for sediment type and organic input), as by host species (90%, see Figure 1). The three panels show non-metric multi-dimensional scaling (NMDS) plots of UniFrac dissimilarity values calculated from estimated relative abundances of symbiont genera across host individuals for ocean basin (on the left), sediment type (middle) and organic input (right). UniFrac stress value: 0.136. For each metadata category, PERMANOVA r^2^ values are indicated within the respective plot; r^2^ values shown in bold were statistically significant. Abbreviations: AO Atlantic Ocean, MS Mediterranean Sea, RS Red Sea, IO Indian Ocean, PO Pacific Ocean, UK unknown, S Sand, CarS Carbonate Sand, CorS Coral Sand, SilS Silicate Sand, M Mud, Si Silt, Al Algae, AlC Algal Crust, AlP Algal Plumbs, C Corals, SG Seagrass, D Detritus.

We detected only minor differences in the symbiont community composition of individuals from the same host species, with highly similar symbiont communities in host individuals from the same sampling site. For the seven host species for which populations could be sampled at multiple sites, with distances between sites ranging from ∼ 245 – 2910 kilometers, symbiont community composition was largely stable, and highly similar compared to differences between host species, including co-occurring species from the same collection site (Tables S1, S3, Fig. S38). However, individuals of the same host species collected at different sampling locations showed higher divergence than individuals from the same geographic location. Hence, slight shifts in the composition of symbiont communities could be a result of increasing intraspecific divergence between hosts from populations separated by hundreds of kilometers or more. Overall, symbiont community composition was highly specific to each host species and consistent across individuals from the same species, with host species as the main factor shaping the composition of symbiont communities in gutless oligochaetes.

### Similarity of symbiont community composition between host species is lost over evolutionary time

Given that symbiont community composition was highly specific to each host species, we asked if very closely related hosts had more similar symbiont communities than more distantly related hosts, a process known as phylosymbiosis^80,81^. Indeed, we found that five out of 21 host sister species had highly similar symbiont communities (Fig. 1, green-blue shading). We confirmed that this observation was linked to host relationships and not a result of random association, by comparing the number of shared nodes between host species and symbiont communities shown in Fig. 1 to 1000 permutations of the symbiont community data (t-test p-value: < 2.2*10^−16^, Fig. S39). Notably, these five clades of host sister species were significantly less diverged than the other 14 host sister species (Wilcoxon signed-rank test p-value: 0.01157, Fig, S40) and more frequently occurred in geographic proximity (Wilcoxon signed-rank test p-value: 7.4*10^−06^, Fig. S40). We then tested for phylosymbiosis across all 63 host species by comparing the topologies of their phylogenetic tree to dendrograms representing the composition of symbiont communities using the Robinson Foulds (RF) metric. The observed congruence between these was 8% higher than expected under random association (RF-to-YuleAvg = 0.9203), indicating the trees were slightly more similar than expected by chance, and did not have a strong phylosymbiosis signal. This topology-based finding was confirmed by a second approach, in which we analyzed the relationship between host phylogenetic distances and symbiont community composition: We found only a very weak linear correlation across the 63 host species analyzed in this study (correlation coefficient of linear model: R^2^=0.04, Fig. S41). Our finding that symbiont communities are only highly similar within host species and between some closely related sister species indicates that partner fidelity dissolves soon after host speciation, with geographic separation amplifying this process.

### To each their own: At the microevolutionary scale, symbiont species are highly similar across host individuals, with considerable within-species symbiont diversity

Our analyses above revealed that the composition of symbiont communities in gutless oligochaetes is stable within host species, but this stability is quickly lost at macroevolutionary scales. To better understand how stable symbiont communities are at the microevolutionary scale, we increased the resolution for the symbionts from 16S rRNA gene analyses to metagenome-assembled genomes (MAGs). We generated 364 medium quality MAGs (estimated genome completeness ≥ 50%, estimated contamination < 10%, Table S5)^82^ for 26 symbiont genera and used their average nucleotide identities (ANIs) for comparative analyses (sequence coverage was not high enough for the remaining 7 of the 33 symbiont genera, Table S6).

In host individuals from the same species, ANIs within each symbiont genus were generally above 95%, the commonly accepted threshold for a bacterial species^83^, while across host species, they were mostly below 95% (Tables S6, S7, Note S5). At a higher resolution of symbiont variability within host species, we observed considerable diversity within symbiont species, with ANIs for the symbionts of individuals from the same host species ranging from 95.03% - 100% (median: 99.72%, mean: 99.31 ± 0.93%). In summary, each host species harbored up to ten symbiont species that were host-specific and distinct from those of other host species, with considerable intraspecific symbiont variability within each host species (Tables S6, S7).

We next asked how similar symbionts were in individuals from the same host species but different populations, to examine if we could detect a decrease in specificity as host populations diverge. We had enough specimens for these analyses from four host species, which were collected at sites separated by 733 - 2911 km (Table S8). In all four host species, genetic differences across individuals from the same population were smaller than those between geographically separated populations for both mtCOI and 28S rRNA, indicating ongoing speciation through isolation by distance (Table S8). However, there was no consistent degree of symbiont divergences in the four host species. For example, in *I. leukodermatus* separated by > 2900 km, symbionts belonged to the same species with ANIs highly similar at ≥ 98%, including the main symbiont *Thiosymbion*. In contrast, in *O. algarvensis* separated by 733 km, symbiont ANIs differed considerably and symbionts, even *Thiosymbion*, clearly belonged to different species (78 - 85%, Table S8). These results show that extensive analyses along a geographic gradient with multiple populations are needed to fully understand the role that isolation by distance plays in gutless oligochaetes and their symbionts.

### Partner fidelity at the symbiont strain level is weak in most host species

To further characterize the processes shaping partner fidelity between gutless oligochaetes and their symbionts at microevolutionary timescales, we compared host mitochondrial COI and nuclear 28S rRNA gene divergences to symbiont ANIs (Fig. 3, Figure S42, Table S9) We analyzed these two genes because they reflect different drivers of partner fidelity. Mitochondria are inherited vertically through the maternal line, and a strong correlation between the genetic distances of a host’s mitochondrial genes and symbiont genes indicates vertical transmission of symbionts^37,84,85^. In comparison to mtCOI, the nuclear 28S rRNA gene evolves much more slowly in most animals, is biparentally inherited, and is a good indicator of host genotype divergence^86,87^. We therefore interpret a strong correlation between genetic distances of host 28S rRNA and symbiont genes as an indication of long-term fidelity, driven by selective recruitment of symbionts via nuclear-encoded host genes such as those involved in immunity.

**Figure 3:**
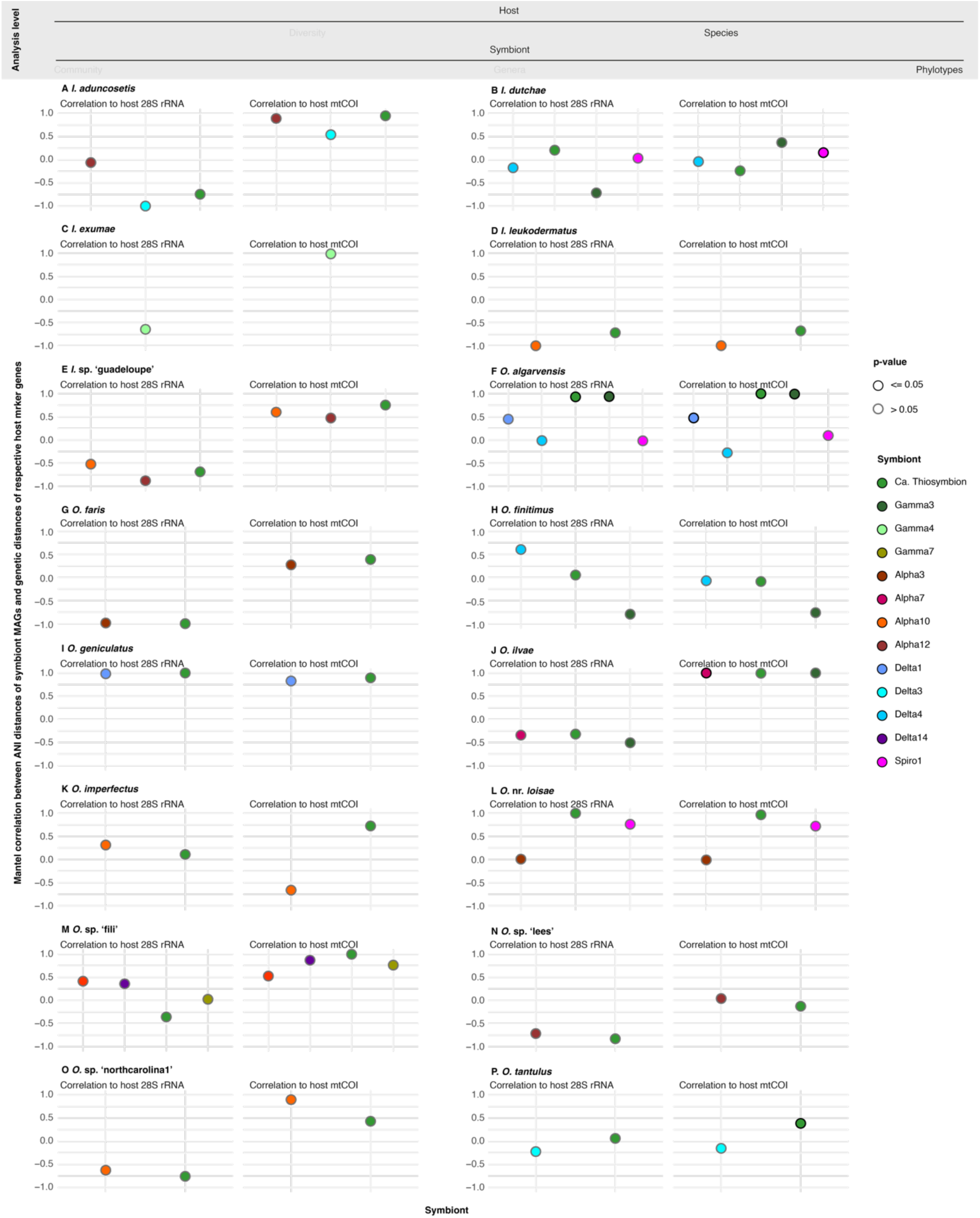
Partner fidelity varies across gutless oligochaete species and is generally stronger for mitochondrial than nuclear genotypes. Each panel shows the correlation values for the indicated host species between symbiont ANIs and the host nuclear gene 28S rRNA (left), and the host mitochondrial gene COI (right). For all data points in this image, MAGs for each symbiont genus were recovered from at least three individuals per host species. Significant correlations (Mantel test; p ≤0.05) are highlighted.

Our analyses revealed that long-term partner fidelity was low in most host species, based on poor correlations between host nuclear 28S rRNA and symbiont ANIs, even for *Thiosymbion* (Fig. 3, Fig. S41; Mantel test R ≥ 0.6 in only 5 out of 16 host species and only for some of a given host’s symbionts; and 7 out of 40 tested host-symbiont pairings.) For host mitochondrial genotypes, fidelity was generally higher than for 28S rRNA (t-test p = 0.004283), but correlations were only strong and positive in 7 out of 12 host species and never for all symbionts of a given host species (Fig. 3, Mantel test R ≥ 0.6 in 8 out of 13 host species, and 14 out of 33 host-symbiont pairings). Unexpectedly, the fidelity of *Thiosymbion* to mitochondrial genotypes was not high in all host species (Fig. 3), despite the symbiont’s role as the host’s main nutritional source and a previous study showing strict mitochondrial fidelity of *Thiosymbion* in *O. algarvensis*^37^ (also confirmed here, Fig. 3). Also unexpected was the high fidelity of some secondary symbionts (Fig. 3), in contrast to the moderate to low fidelity observed in the secondary symbionts of *O. algarvensis*^37^. These results underscore the importance of avoiding the assumption that findings from one symbiont or host can be generalized and applied to other related symbionts or hosts, even very closely related ones.

### At macroevolutionary scales, partner fidelity is low in most symbiont genera

To gain insights into how stable host-symbiont partnerships are over macroevolutionary timescales, we extended our analyses from symbiont fidelity within host species to patterns across host species. Specifically, we asked whether fidelity varies across host species for a given symbiont genus, that is, whether some symbiont genera have co-diverged more closely with gutless oligochaetes than others. To answer this question, we used MAGs from genera that were present in at least four host species, leaving us with 12 symbiont genera for testing correlations between symbiont and host genetic distances (for the symbiont MAGs we used ANIs, for the host 28S rRNA and mtCOI genes; Fig. 4). In all symbiont genera, ANIs were positively correlated with genetic distances of both host genes, indicating a certain degree of partner stability and fidelity over macroevolutionary time (Fig. 4, Mantel test R 0.32 to 0.93). However, strong correlations between host and symbiont genetic distances (Mantel test R >0.6), were observed in only six symbiont genera for 28S rRNA, and in nine symbiont genera for mtCOI (Fig. 4, Fig. S43). Most symbiont genera, including *Thiosymbion*, had low correlation coefficients, with many host - symbiont pairings deviating from the linear regression line (dotted lines in Fig. 4; e.g. *Thiosymbion* or Alpha12). While closely related hosts often had closely related symbionts, there was substantial variability with increasing genetic distances between hosts (Fig. 4). These outliers are a clear indication of horizontal symbiont acquisition, either through host switching or uptake from the environment. While horizontal symbiont acquisition was shown previously for *Thiosymbion* in 22 gutless oligochaete species based on 16S rRNA analyses^67^, we show here that disruption of codivergence is common across many symbiont genera of gutless oligochaetes.

**Figure 4:**
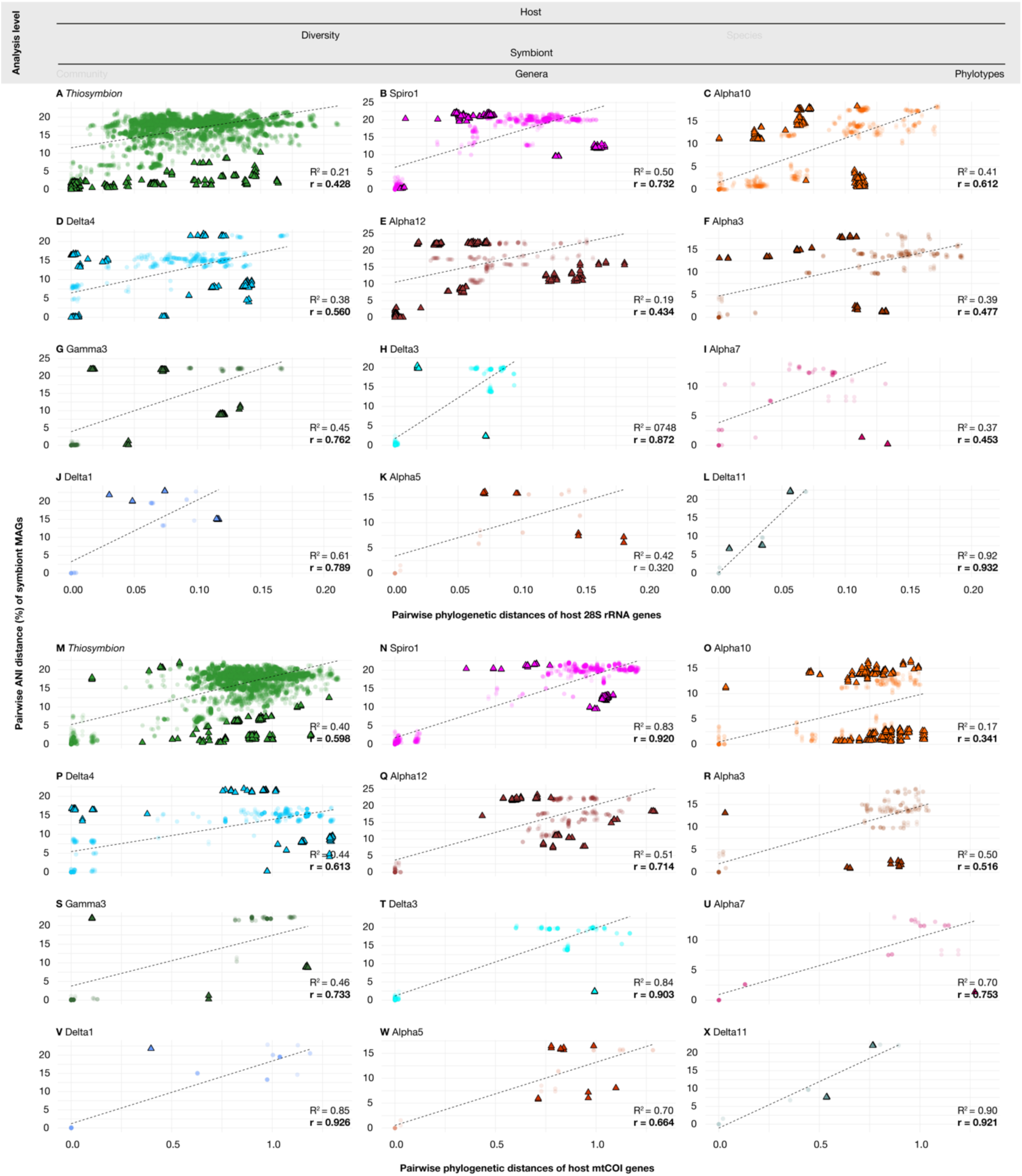
Partner fidelity varies across symbiont genera. Each panel shows the pairwise ANI dissimilarity from MAGs of a given symbiont genus with the corresponding pairwise genetic dissimilarity of host marker genes. **A–L:** ANI distances plotted against 28S rRNA distances of the host. **M–X:** ANI distances plotted against mtCOI distances of the host. Dotted lines represent linear regression lines, with R² values indicating the proportion of variance explained by the regression and r values representing correlation strength from Mantel tests. Bold r values denote statistically significant correlations (p ≤ 0.05). Two types of data points are shown based on their perpendicular distance to the regression line: solid triangles show the 50% of host-symbiont pairings farthest from the regression, transparent circles the 50% closest to the regression, to better visualize outliers from host-symbiont codivergence.

In some symbiont genera, codivergence of symbiont ANIs with host mtCOI was considerably higher than with 28S rRNA (e.g. Alpha12 or Alpha5, Fig. 4), indicating that in these, maternal transmission is the main mode of acquisition. Conversely, in symbiont genera in which codivergence was higher with 28S rRNA than with mtCOI, such as Alpha10, symbionts could be transmitted paternally or acquired horizontally from the environment or co-occurring host species, with these de novo acquisitions likely shaped by nuclear-encoded selection factors.

### *De novo* acquisition, host switching and loss of symbionts drive variability and novelty in symbiotic communities on macroevolutionary scales

To understand how long symbiont genera have been associated with gutless oligochaetes, and identify gains and losses over time, we estimated the ages of gutless oligochaetes and their symbiotic genera using molecular clock models, and ancestral state reconstruction (Fig. 5, Table S10). Our analyses estimated the age for the radiation of gutless oligochaete at 159 Ma, much older than previous analyses of 50 Ma^64^. This is likely due to our broader sampling of gutless oligochaete species, as well as the inclusion of closely related non-symbiotic oligochaetes. Based on our age estimates for the symbiont genera, *Thiosymbion* was acquired first, before the diversification of gutless oligochaetes. Loss of *Thiosymbion* occurred in only a few species, namely *I. exumae*, as shown previously in seven individuals collected in 1999^56^ and confirmed in this study for three individuals collected 14 years later; *O. sp.* ‘other’ (n=2/2, this study); and in *O. sp.* ‘belize2’ (n=2/3, this study). It is therefore reasonable to assume that the acquisition of *Thiosymbion* as a primary producer of nutrition via chemosynthesis was a key innovation and major driver of gutless oligochaete diversification. As to vice-versa - whether the establishment of symbioses with gutless oligochaetes drove the diversification of *Thiosymbion* – this remains unclear because these bacteria are also symbionts of marine nematodes^67^, and it is unknown which host group they initially associated with. The estimated median age of *Thiosymbion* of about 415 Ma and their ectosymbiotic association with stilbonematinid nematodes, a lifestyle thought to predate endosymbiosis, suggest that *Thiosymbion* diversified over a longer evolutionary time before establishing associations with gutless oligochaetes.

**Figure 5:**
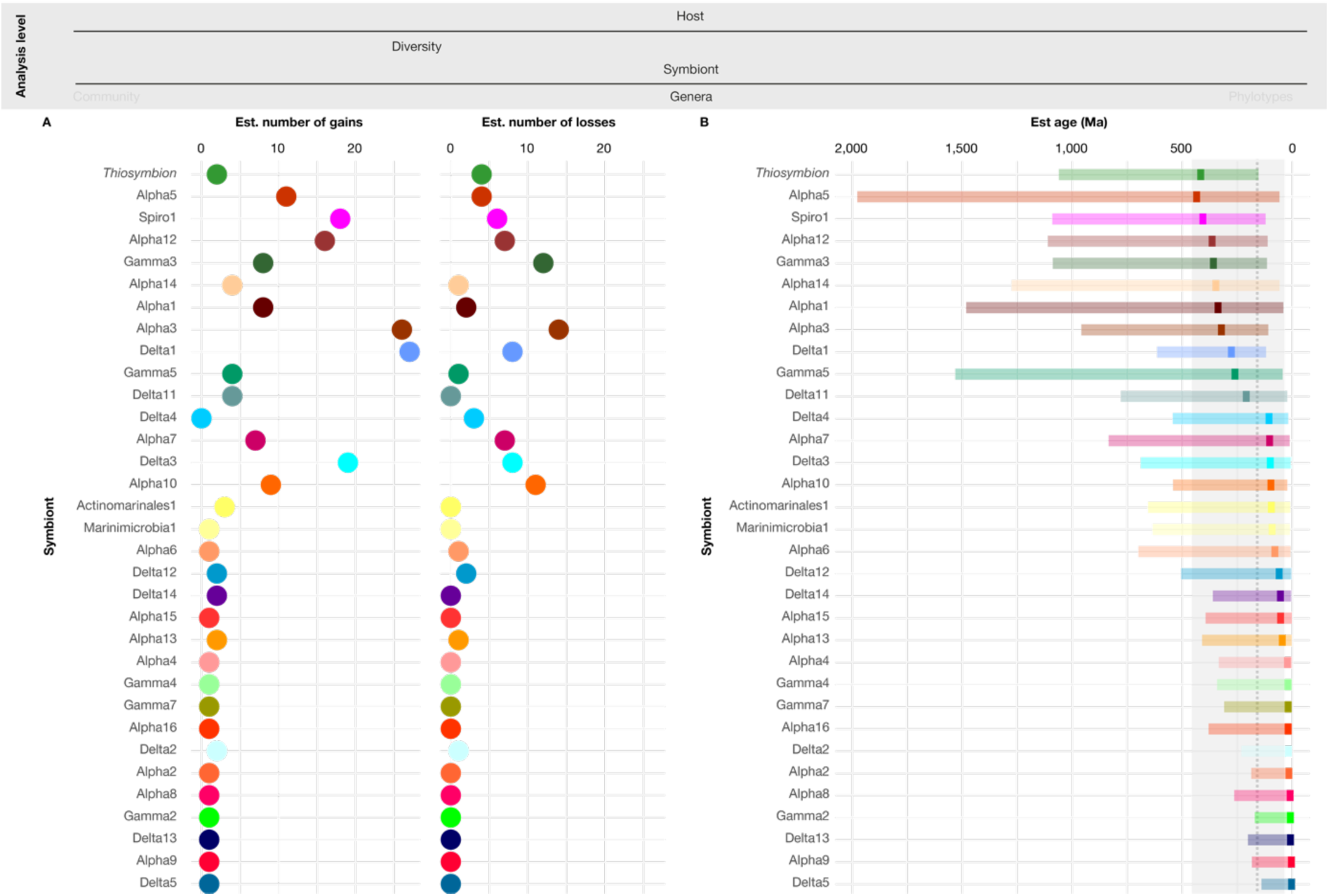
The primary symbiont *Thiosymbion* was acquired only once, while secondary symbiont genera were likely gained and lost several times. **A)** Shows estimated number of acquisitions (left panel) and losses (right panel) of each symbiont genus based on ancestral state reconstructions using **PastML**^88^. **B**) Shows the estimated divergence times of symbiont genera relative to the gutless oligochaete clade (grey shading) based on a Bayesian analysis framework with a relaxed log normal clock model. Symbiont genera are listed on the y-axes, sorted by their estimated median ages (top: older, bottom: younger).

In the secondary symbionts, that is all other symbiont genera besides *Thiosymbion*, we observed two patterns for gains (Fig. 5, Table S10). Widespread symbiont genera associated with several, often distantly related host species were likely acquired multiple times throughout the diversification of gutless oligochaetes in convergent evolution, in some cases ten times or more, such as the Alpha3, Alpha10 and Gamma3 symbionts (Fig. 1 and Fig. 5). The estimated ages of these genera were relatively old. In contrast, the secondary symbionts confined to a single host clade were relatively younger (Fig. 1 and Fig. 5) and likely acquired recently and only once by the last common ancestor of the host clade. Based on our phylogenetic analyses of symbiont genera and their closest relatives showing that most symbiont genera were not intermixed with environmental sequences, the majority of these repeated acquisitions likely happened via acquisition from co-occurring host species, rather than acquisition from the surrounding environment (phylogenetic analyses of 16S rRNA sequences using data generated in this study and from public repositories; Fig. S2-34, Note S2).

In addition to repeated convergent events of symbiont gains, losses were also widespread in secondary symbiont genera, but less common than symbiont gains (187 predicted gains vs. 93 losses; 2.01 gains/losses ratio across 33 symbiont genera; Fig. 5, Table S10). A higher ratio of symbiont gains over losses indicates that the number of symbiont genera has increased over time in gutless oligochaetes, leading to an increase in the diversity and complexity of their symbiotic communities.

## Conclusions

In this study, we explored the specificity, genetic variability and evolution of symbiont communities in gutless oligochaetes, examining both microevolutionary dynamics within individual host and symbiont species, as well as macroevolutionary processes across both host species and symbiont genera. By integrating these two scales in both the host and symbionts, we gained insights into the evolutionary processes that shape these diverse multi-partner in which a clade of animal hosts has become obligately dependent on their bacterial symbionts for their nutrition and waste removal.

Our analyses showed that host species had by far the highest explanatory power for differences in symbiont community composition at 90%, with ocean basins and environmental settings playing much more minor roles (Fig. 2). Given the obligate dependency of gutless oligochaetes on their symbionts, it is not surprising that the composition of symbiont communities within host species is not random, but rather highly consistent. This specificity is driven by vertical transmission of symbionts from parents to offspring through the smearing of eggs with symbionts as the eggs exit the mother^68–71^. The eggs are then fertilized with sperm stored in the maternal spermathecae, and a cocoon formed around the embryo and symbionts, which is then deposited in the surrounding sediment ^68–71^. This form of vertical transmission offers the advantage of consistent transmission of parental symbionts, while also providing opportunities for bacteria from the environment or co-occurring hosts to invade before the cocoon becomes too hard to penetrate. Indeed, in *O. algarvensis*, the only host species in which the fidelity of its symbiont species was analyzed in detail, already within a single host population, fidelity was not strict across the symbiotic community, indicating non-maternal acquisition of bacterial symbionts^37^. Notably, however, the newly acquired bacteria were clearly limited to strains of the same bacterial species^37^. This pattern now holds true for all gutless oligochaetes: Our 16S rRNA sequence analyses showed strong specificity between hosts and their symbiotic species, indicating a highly selective process in which hosts and their symbionts interact with each other to enable colonization of the developing embryo and persistence throughout the lifetime of the host. This specificity, however, is much less pronounced at the symbiont strain level, as visible in the relatively low levels of partner fidelity in our comparisons of symbiont ANIs to host genetic distances (Figs. 3 and 4). Similar patterns of species-level symbiont specificity combined with strain-level variability are increasingly being observed in both marine and terrestrial hosts, thanks to advances in both high-throughput and long-read sequencing that now allow researchers to detect intraspecific bacterial diversity, particularly in non-cultivable symbionts^30,89–91^.

As gutless oligochaete species diverge, the strict selection of symbionts at the species level weakens (Fig. 1). New symbiont species or even genera and families are acquired, either from other hosts or the environment, and existing symbionts are displaced or lost (Fig. 5). Several factors could explain this remarkable flexibility in acquiring novel symbionts. First, the consistent association of gutless oligochaetes with their dominant primary symbiont *Thiosymbion*, which was likely acquired when these hosts first evolved and is present in nearly all recent hosts, may fulfil enough critical needs of the hosts to allow considerable flexibility in the acquisition of secondary symbionts. Second, the extracellular location of the symbionts below the worm’s cuticle and above their epidermal cells may require less stringent specificity than an intracellular symbiosis. Third, during the deposition of eggs in the sediment, there are numerous opportunities for bacteria from the environment to enter the developing cocoon. Fourth, the displacement of established symbionts by novel lineages may prevent symbiont cheating as well as genomic decay^92–94^. Finally, and importantly, there is a strong selective advantage in acquiring novel symbionts. These generate functional diversification, by providing hosts with more, potentially complementary or syntrophic, metabolic pathways for energy conservation, carbon fixation, respiration, resource partitioning and recycling. This metabolic diversity is likely highly adaptive, particularly in environments where resources are limited, as shown for *O. algarvensis* and its five co-occurring symbionts^63,66,95^.

In summary, our study shows that the symbiotic communities of gutless oligochaetes remain highly stable at microevolutionary scales, yet shift rapidly over short macroevolutionary timescales as host species diverge. Similar processes have been observed in the few studies that integrate both micro- and macroevolutionary timescales in multipartite host-microbe associations. However, these studies have focused on gut microbiomes, where microbes are not embedded within host tissues, and host diet and ecology over the lifespan of the animal play major roles^96,97^. In contrast, gutless oligochaetes acquire their symbionts only during a narrow window associated with sexual reproduction and these bacteria must persist throughout embryonic and juvenile development. The dynamic balance in gutless oligochaete symbioses between evolutionary stability over short time scales and consistent remodeling of their symbiont communities with repeated acquisitions of novel symbiont lineages over longer time scales is thus distinctive, and may be a key factor underlying the evolutionary success of this widespread and diverse host group.

## Material and methods

### Sample collection, processing and metagenomic sequencing

231 individuals of gutless and 10 individuals of gut-bearing oligochaetes were sampled between 1991 and 2018 at sites around the world (for overview see Table S1). Individual specimens were rinsed and either flash-frozen in liquid nitrogen and stored at −80°C or fixed in RNAlater (Thermo Fisher Scientific, Waltham, MA, USA) and stored at 4°C or −20°C. DNA was extracted from single worm individuals with the DNeasy Blood & Tissue Kit (Qiagen, Hilden, Germany) according to the manufacturer’s instructions. Library construction, quality control and sequencing were performed at the Department of Energy Joint Genome Institute (Berkeley, California, USA) and the Max Planck Genome Centre (Cologne, Germany). Information on library preparation and sequencing details are listed in Table S11. Some samples were sequenced twice to generate a higher number of reads. In these cases, resulting reads from both sequencing runs were combined for further analyses. Two samples were extracted twice using different library preparation methods and individually sequenced. The resulting sequences were also pooled for further analyses.

### Assembly of host marker genes

28S rRNA and mtCOI genes of all specimens were assembled by mapping the metagenomic reads to respective databases using bbmap v38.34 (https://sourceforge.net/projects/bbmap/). For the 28S rRNA, we used the SILVA database v138^98,99^. Mapped reads were assembled using SPAdes v3.11.0 setting k-mer sizes to 99, 111 and 127 bp^100^. Final 28S rRNA gene sequences were predicted from the assembled sequences using barrnap v0.9 (https://github.com/tseemann/barrnap). MtCOI genes were assembled by adapting the phyloFlash pipeline to operate on a custom mtCOI reference database and predict mtCOI genes from assembled sequences^101^. In the few cases in which more than one mtCOI gene was found in a sequencing library, we used the most abundant mtCOI sequence based on read coverage. If no dominant and phylogenetically coherent mtCOI sequence that corresponded to the 28S rRNA’s phylogenetic results could be identified, the mtCOIs for these libraries were excluded from strain level analyses.

### Identification of host taxa

Host taxa were defined based on mtCOI gene phylogenies (see below), which included the published gene sequences of gutless oligochaetes previously identified using both morphological and molecular traits. Specimens that could not be assigned to published species based on morphological or molecular data were treated as new taxa and assigned provisional names with consecutive numbers and the sampling location. For the gut-bearing oligochaetes used in this study, the presence of a gut was determined based on morphological observations.

### Host marker gene phylogenies

28S rRNA and mtCOI gene sequences were aligned using MAFFT-XINS-i and MAFFT-LINS-i v7.407 respectively^102–104^. The mtCOI alignment was manually trimmed in Geneious v11.1.5 and bases 40-695 were kept (https://www.geneious.com). Maximum-likelihood based phylogenies were calculated using IQ-TREE, including automatic selection of the best suited model and generation of 100 none-parametric bootstrap replicates. The sequences of one gut-bearing oligochaete specimen (*Phallodrilinae* gen. sp. ‘strang’) were used to root the phylogenies in iTOL^105^. To compare the 28S rRNA and the mtCOI gene-based host phylogenies, we calculated the number of shared nodes between the maximum-likelihood trees calculated for each of the genes using the R package ‘ape’^106^.

### Symbiont clade definition and quantification

16S rRNA genes were assembled from the metagenomic libraries of gutless oligochaetes via phyloFlash using the –all option and in addition specifying the read length. For subsequent analyses, we only considered sequences that were i) assembled with SPAdes, ii) longer than 1000 bp and iii) did not contain more than 20 ambiguous bases. The resulting sequences were clustered at 95% sequence similarity using usearch v10.0.240^107^. We used the SINA search and classify algorithm to add the 16S rRNA gene sequences of close relatives from the SILVA database v132 that shared at least 90% sequence similarity for each of our assembled symbiont sequences^108^. All assembled sequences and the SILVA database hits were aligned using MAFFT-LINS-ii and a phylogenetic tree was calculated from the resulting alignment using FastTree v2.1.1^109^. We mapped the 95% clusters to this tree and manually merged monophyletic clades that consisted of several of the 95% clusters into single symbiont clades. We analyzed the prevalence of all phylogenetically defined symbiont clades across the gutless oligochaete metagenomic libraries. We excluded clades that had distribution patterns that suggested they were contaminations or spurious associations (Note S1). The abundances of the remaining clades (symbiont genera from here on) were quantified across all metagenomic libraries using EMIRGE v.0.61.1 following the standard workflow for custom EMIRGE databases^110^.

### Phylogeny of all symbionts and their relatives

All sequences included in the symbiont genera defined above were used to obtain sequences from closely related bacteria from the SILVA and the RefSeq public databases^111^. For SILVA, we used the SINA search and classify algorithm to obtain up to 10 relatives for each sequence that shared at least 99% and 95% sequence similarity for each of our input sequences. In addition, we also screened the RefSeq database using BLAST implemented in Geneious v11.1.5 to obtain the ten most similar 16S rRNA genes^112^. Duplicated sequences were removed from the collection of sequences of the symbionts’ relatives. In addition, we included the 16S rRNA gene sequence of *Crenarchaeotal* sp. clone JP41 (NCBI accession: L25301.1) as outgroup. Sequences were aligned with MAFFT-XINS-i. Maximum-likelihood based phylogenies were calculated using IQ-TREE, including automatic selection of the best suited model and generation of 100 none-parametric bootstrap replicates^121–123^. We also calculated a separate tree for each symbiont genus using sequences assembled in this study and reference sequences from SILVA and RefSeq. For SILVA, we used the SINA search and classify algorithm to obtain up to 10 relatives for each sequence that shared at least 99%, 95% and 90% sequence similarity for each of our input sequences. In addition, we also screened the RefSeq database using BLAST implemented in Geneious v11.1.5 to obtain the ten most similar 16S rRNA genes. Duplicated sequences were removed from the collection of sequences of the symbionts’ relatives. In addition, we included the 16S rRNA gene sequence of *Crenarchaeotal* sp. clone JP41 (NCBI accession: L25301.1) as outgroup. Sequences were aligned with MAFFT-XINS-i. Maximum-likelihood based phylogenies were calculated using IQ-TREE, including automatic selection of the best suited model and generation of 100 none-parametric bootstrap replicates.

### Analyses and plotting of symbiont community composition

The analyses of symbiont community composition were performed in R v4.4.1 unless differently stated. During the analyses, the following packages were used: phyloseq^113^, ape, vegan (https://github.com/vegandevs/vegan), plyr^114^, MASS^115^, gdata (https://cran.r-project.org/web/packages/gdata/index.html), reshape2 (https://github.com/hadley/reshape), forcats (https://github.com/robjhyndman/forecast), igraph (https://github.com/igraph/rigraph), Hmisc (https://github.com/harrelfe/Hmisc/), optparse (https://github.com/trevorld/r-optparse), data.table (https://github.com/Rdatatable/data.table), ade4 (https://github.com/sdray/ade4), tidyverse^116^ and spa (https://github.com/markvanderloo/rspa). Plots were generated using ggplot2 from the tidyverse package, gridExtra (https://cran.r-project.org/web/packages/gridExtra/index.html), ggpubr (https://cran.r-project.org/web/packages/ggpubr/index.html), maps (https://www.rdocumentation.org/packages/maps), mapdata (https://www.rdocumentation.org/packages/mapdata), and patchwork (https://github.com/thomasp85/patchwork).

### Community composition analyses

The similarity between symbiont communities of host individuals were calculated based on the abundance patterns of the symbiont genera and the symbiont 16S rRNA gene phylogeny using the UniFrac metric as implemented in the phyloseq package in R. We tested for parameters that could explain differences in symbiont community composition between individuals using PERMANOVA and the Mantel test^117,118^. These parameters included host species, geographic distance, ocean, organic input and sediment type (Table S1). All factors except for geographical distances were treated as categorical data and analyzed using PERMANOVA. Geographical distance was treated as the correlation between the UniFrac distances and actual geographic distances and analyzed using the Mantel test.

Co-occurrence patterns of symbiont genera were analyzed using Spearman’s correlations and were corrected using the Benjamini-Hochberg standard false discovery rate correction. Networks of significant co-occurrences were generated using igraph^119^.

### Phylosymbiosis

UniFrac distances on the average symbiont abundances per host species were transformed into a dendrogram using hierarchical clustering. The congruences between the 28S rRNA gene-based host tree or and the symbiont community UniFrac dendrogram was assessed using the Robinson-Foulds metric, implemented in TreeCmp v1.0-b291^120,121^.The relation between host phylogenetic distances and the symbiont community composition distances was analyzed using linear regression and the Mantel test.

### Generation of symbiont MAGs and calculation of ANI values

Symbiont MAGs were generated using a custom pipeline that combines trimming and filtering of raw reads, metagenomic assembly and binning. The exact pipeline is shared under: https://github.com/amankowski/MG-processing_from-reads-to-bins/tree/main. Bin quality was examined by genome completeness and contamination (checkM v2)^122^, the presence of a 16S rRNA gene (barrnap v0.9, https://github.com/tseemann/barrnap) and the number of amino acids that had at least one tRNA encoded in the genome as a second proxy for genome completeness (ARAGORN v1.2.38)^123^. Taxonomy was assigned to bins that were of at least medium quality according to MIMAG standards^100^ (> 50% complete and showed < 10% contamination, GTDBtk v1.3.0, GTDB r95)^124–126^. In addition, we also determined the GTDB taxonomy of previously produced symbiont bins which were assigned to a genus based on 16S rRNA phylogenies. We assigned a symbiont genus to each bin based on the shared GTDBtk taxonomy with our internal reference bins. If multiple bins were present per library and symbiont genus, we selected the best one based on genome completeness and contamination estimates, the presence of 16S rRNA, and the number of amino acids with at least one tRNA encoded. We then removed contigs that were shared by at least two bins from the same library. The final bins were checked again with checkM v2 and the genome statistics of these final bins are reported in Table S5. ANI values between all MAGs from the same symbiont genus were calculated using FastANI v1.33^83^ and the resulting distances were used for further correlation analyses.

### Correlation between host and symbiont phylogenetic distances

For all hosts with member sequences of a given symbiont genus we calculated pairwise phylogenetic distances of the hosts’ 28S rRNA or the mtCOI genes R’s cophenetic function. We analyzed the correlation between the host genetic distance and symbiont ANI distances using linear regression and the Mantel test.

### Estimates of divergence times for host and symbiont clades and reconstruction of ancestral states of symbiont association patterns

For the estimation of host divergence times, we used a Bayesian phylogenetic framework and a relaxed molecular clock model. We constructed a matrix of eight 28S rRNA gene sequences of gutless host and eight publicly available 28S rRNA gene sequences of other Oligochaeta and one representative of the Polychaeta. The oligochaete representatives were selected to i) cover a broad diversity of the phylum and ii) to include the following calibration points for our molecular clock model: the last common ancestor of the Goniadidae (323 Ma), the last common ancestor of the Hormogastridae (82 ± 15 Ma), the divergence between Hirudinea and Lumbriculidae (201 Ma) and the last common ancestor of the Phyllodocida (485 ± 1.9 Ma)^127,128^. The polychaete sequence was included to root the tree and to include the last common ancestor of all Annelida (510 ± 10 Ma) as additional calibration point. All calibration points were considered as uniform priors. All sequences were aligned using mafft-linsi and the time calibrated tree was calculated in BEAST v2.6.3 using the GTR+G+I model and the relaxed log normal clock model^129^. Besides the mentioned priors for time calibration, we set Alpha and Beta values of birtRate.Y.t prior to 0.001 and 1,000, respectively. We also defined priors that considered the oligochaetes and the gutless oligochaetes as monophyletic groups. The chain length was set to 100,000,000 and the sample frequency was set to 1,000. All estimated parameters were controlled to show an ESS > 200 in Tracer. We used TreeAnnotator to calculate the node ages (i.e., mean heights) after setting a burn-in of 25 %.

We used the same approach for the host to estimate the divergence times for symbiont genera based on a subset of our symbiont sequence matrix that combined 2-3 symbiont 16S rRNA gene sequences per symbiont genus with 50 type strain sequences from RefSeq databases. Type strains of the *Chromatiaceae* were used to include their previously published divergence estimate as calibration point for the symbiont analysis^130^. In addition, we included the 16S rRNA gene sequence of *Crenarchaeotal* sp. clone JP41 (NCBI accession: L25301.1) as an outgroup. The time calibrated tree was calculated in BEAST v2.6.3, using the GTR+G+I model and the relaxed log normal clock model. The divergence time of the *Chromatiaceae* was considered as an exponential prior with a mean value of 0.1 and an offset of 1.64. We additionally constrained the analyses by setting a uniform prior from 3.5-4.5 billion years for the whole dataset to account for the maximum age of life on earth. Additional priors were used to define monophyletic clades for all bacteria, the Delta1, Delta4 and Delta12 genera as well the combined Delta4-Delta12 clade that were observed in the previous phylogenetic analyses of the symbiont genera. We set Alpha and Beta values of birtRate.Y.t prior to 0.001 and 1,000, respectively. We ran 4 parallel chains, setting the chain length to 500,000,000 generation and the tree sample frequency was set to 1,000. All estimated parameters were controlled to show an ESS > 200 in Tracer. Afterwards, trees were downsampled from 500,000 to 100,000 using LogCombiner and we used TreeAnnotator to calculate the node ages (i.e., mean heights) after setting a burn-in of 25 %. Ancestral states of symbiont presence/absence patterns were performed using a maximum likelihood based last common ancestor analysis with PastML (v1.9.45), using the MAP prediction method^88^.

## Data and script availability

Raw metagenomic sequences produced by the DOE JGI are available at the JGI Genome Portal under GOLD Study ID Gs0095504. Raw metagenomic sequences produced by the Max Planck Genome center are deposited in the European Nucleotide Archive (ENA) under accession ID PRJEB94555 and will be made available upon publication of the manuscript. Additionally, we deposited marker gene sequences, phylogenetic trees and MAGs generated in this study (https://doi.org/10.5281/zenodo.16568832). The data can be shared with the reviewers upon request and will be made publicly available upon publication of the manuscript. The scripts and data for analyzing symbiont community composition and phylogenetic correlations are available under https://github.com/amankowski/Mankowskietal_symbiont-diversity-in-GOs.

## Acknowledgments

We are thankful for sample collections and field assistance by Alexander Gruhl, Anna Ansebo, Anna Blazejak, Anne-Christin Kreutzmann, Christian Lott, Claudia Bergin, Dolma M. Michellod, Emilia M. Sogin, Erica Mejlon, Erich Mueller, Falk Warnecke, Fred Wells, Jörg Ott, Judith Zimmermann, Katrine Worsaae, Ken Halanych, K. B. Brandon Seah, Lisa Matamoros, Lena Gustavsson, Mario P. Schimak, Michael Hadfield, Miriam Sadowski, Miriam Weber, Olav Giere, Oliver Jäckle, Nicholas Bekkouche, Olivier Gros, Pamela Reid, Philippe Bouchet, Pierre De Wit, Ramon Rosello-Mora, Silke Wetzel, Silvia Bulgheresi, Stefan Sommer, and Tina Enders. In addition, we would like to thank the crew of the Meteor research cruise M92 and the Sonne research cruise SO147, as well as the Carrie Bow Cay Laboratory, the Heron Island Research Station, the HYDRA Institute Elba, the Mediterranean Institute for Advanced Studies, the Lee Stocking Island Research Station, the Little Darby Island Research Station, the Lizard Island Research Station, and the Okinawa Institute of Science and Technology and their staff for supporting our sampling campaigns. This work was supported by the Max Planck Society, a Moore Foundation Marine Microbial Initiative Investigator Award to ND (Grant GBMF3811), the MARUM Cluster of Excellence ‘The Ocean Floor’ (Deutsche Forschungsgemeinschaft (German Research Foundation) under Germany’s Excellence Strategy - EXC-2077 – 39074603), a U.S. National Science Foundation award to MK (grant IOS 2426305), the USDA National Institute of Food and Agriculture Hatch project 7002782 awarded to MK, and a Heisenberg Grant from the DFG (GR 5028/1-1) and a Marie-Curie Intra-European Fellowship PIEF-GA-2011-301027 CARISYM awarded to HRGV. The work (proposal: 10.46936/10.25585/60000875) conducted by the U.S. Department of Energy Joint Genome Institute (https://ror.org/04xm1d337), a DOE Office of Science User Facility, is supported by the Office of Science of the U.S. Department of Energy operated under Contract No. DE-AC02-05CH11231. This work is contribution XXX from the Carrie Bow Cay Laboratory, Caribbean Coral Reef Ecosystem Program, National Museum of History, Washington DC.

## Author contributions

AM, MK, HRGV, JW and ND conceived the study and AM designed the workflows with input from HRGV and JW. AM, MK, CÉ, NL, YS, JMV, BH, CW, TW, JW and HRGV acquired specimens and generated metagenomic data. AM analyzed the data. AM, HRGV, MK and ND interpreted the results. AM drafted the manuscript and AM, HRGV and ND edited the manuscript. All authors provided revisions.

## Literature

1. Sonnenburg, J. L. & Bäckhed, F. Diet-microbiota interactions as moderators of human metabolism. Nature 535, 56–64 (2016).

2. Gruber-Vodicka, H. R., et al. *Paracatenula*, an ancient symbiosis between thiotrophic *Alphaproteobacteria* and catenulid flatworms. Proc. Natl. Acad. Sci. U.S.A. 108, 12078–12083 (2011).

3. Cavanaugh, C. M., Gardiner, S. L., Jones, M. L., Jannasch, H. W. & Waterbury, J. B. Prokaryotic cells in the hydrothermal vent tube worm *Riftia pachyptila* Jones: Possible chemoautotrophic symbionts. Science 213, 340–342 (1981).

4. Aylward, F. O. et al. Convergent bacterial microbiotas in the fungal agricultural systems of insects. mBio 5, (2014).

5. McCutcheon, J. P. & Von Dohlen, C. D. An interdependent metabolic patchwork in the nested symbiosis of mealybugs. Curr. Biol. 21, 1366–1372 (2011).

6. Bell-Roberts, L., Douglas, A. E. & Werner, G. D. A. Match and mismatch between dietary switches and microbial partners in plant sap-feeding insects. Proc. R. Soc. B 286, (2019).

7. Brune, A. Symbiotic digestion of lignocellulose in termite guts. Nat. Rev. Microbiol. 12, 168–180 (2014).

8. Parish, A. J., Rice, D. W., Tanquary, V. M., Tennessen, J. M. & Newton, I. L. G. Honey bee symbiont buffers larvae against nutritional stress and supplements lysine. ISME J. 16, 2160–2168 (2022).

9. Reis, F. et al. Bacterial symbionts support larval sap feeding and adult folivory in (semi-)aquatic reed beetles. Nat. Commun. 11, (2020).

10. Buysse, M. & Duron, O. Evidence that microbes identified as tick-borne pathogens are nutritional endosymbionts. Cell 184, 2259–2260 (2021).

11. Russell, J. A. et al. Bacterial gut symbionts are tightly linked with the evolution of herbivory in ants. Proc. Natl. Acad. Sci. U.S.A. 106, 21236–21241 (2009).

12. Ott, J., Bright, M. & Bulgheresi, S. Symbioses between Marine Nematodes and Sulfur-oxidizing Chemoautotrophic Bacteria. Symbiosis 36, 103–126 (2004).

13. Petersen, J. M. et al. Dual symbiosis of the vent shrimp *Rimicaris exoculata* with filamentous *gamma*- and *epsilonproteobacteria* at four Mid-Atlantic Ridge hydrothermal vent fields. Environ. Microbiol. 12, 2204–2218 (2010).

14. Petersen, J. M. et al. Chemosynthetic symbionts of marine invertebrate animals are capable of nitrogen fixation. Nat. Microbiol. 2, 16195 (2016).

15. McFall-Ngai, M. J. The importance of microbes in animal development: Lessons from the squid-vibrio symbiosis. Annu. Rev. Microbiol., 68, 177–194 (2014).

16. McAnulty, S. J. et al. ‘Failure To Launch’: Development of a Reproductive Organ Linked to Symbiotic Bacteria. mBio 14, (2023).

17. Habetha, M., Anton-Erxleben, F., Neumann, K. & Bosch, T. C. The *Hydra viridis*/*Chlorell*a symbiosis. Growth and sexual differentiation in polyps without symbionts. Zoology 106, 101–108 (2003).

18. Little, A. F., Van Oppen, M. J. H. & Willis, B. L. Flexibility in algal endosymbioses shapes growth in reef corals. Science 304, 1492–1494 (2004).

19. Kerwin, A. H. et al. Shielding the next generation: Symbiotic bacteria from a reproductive organ protect bobtail squid eggs from fungal fouling. mBio 10, (2019).

20. Kaltenpoth, M., Göttler, W., Herzner, G. & Strohm, E. Symbiotic bacteria protect wasp larvae from fungal infestation. Curr. Biol. 15, 475–479 (2005).

21. Jones, B. W. & Nishiguchi, M. K. Counterillumination in the Hawaiian bobtail squid, *Euprymna scolopes* Berry (*Mollusca: Cephalopoda*). Mar. Biol. 144, 1151–1155 (2004).

22. Jäckle, O. et al. Chemosynthetic symbiont with a drastically reduced genome serves as primary energy storage in the marine flatworm *Paracatenula*. Proc. Natl. Acad. Sci. U.S.A 116, 8505 LP –8514 (2019).

23. De Oliveira, A. L., Mitchell, J., Girguis, P. & Bright, M. Novel Insights on Obligate Symbiont Lifestyle and Adaptation to Chemosynthetic Environment as Revealed by the Giant Tubeworm Genome. Mol. Biol. Evol. 39, (2022).

24. Leigh, E. G. The evolution of mutualism. J. Evol. Biol. 23, 2507–2528 (2010).

25. Archetti, M. et al. Economic game theory for mutualism and cooperation. Ecol. Lett. 14, 1300–1312 (2011).

26. Sachs, J. L., Mueller, U. G., Wilcox, T. P. & Bull, J. J. The Evolution of Cooperation. Q. Rev. Biol. 79, 135–160 (2004).

27. Funk, D. J., Helbling, L., Wernegreen, J. J. & Moran, N. A. Intraspecific phylogenetic congruence among multiple symbiont genomes. Proc. R. Soc. B 267, 2517–2521 (2000).

28. Liu, L., Huang, X., Zhang, R., Jiang, L. & Qiao, G. Phylogenetic congruence between *Mollitrichosiphum* (*Aphididae: Greenideinae*) and *Buchnera* indicates insect-bacteria parallel evolution. Syst. Entomol. 38, 81–92 (2013).

29. Hurtado, L. A., Mateos, M., Lutz, R. A. & Vrijenhoek, R. C. Coupling of bacterial endosymbiont and host mitochondrial genomes in the hydrothermal vent clam *Calyptogena magnifica*. Appl. Environ. Microbiol. 69, 2058–2064 (2003).

30. Ansorge, R. et al. Functional diversity enables multiple symbiont strains to coexist in deep-sea mussels. Nat. Microbiol. 4, 2487–2497 (2019).

31. Petersen, J. M., Wentrup, C., Verna, C., Knittel, K. & Dubilier, N. Origins and Evolutionary Flexibility of Chemosynthetic Symbionts From Deep-Sea Animals. Biol. Bull. 223, 123–137 (2012).

32. Ishii, Y., Matsuura, Y., Kakizawa, S., Nikoh, N. & Fukatsua, T. Diversity of bacterial endosymbionts associated with macrosteles leafhoppers vectoring phytopathogenic phytoplasmas. Appl. Environ. Microbiol. 79, 5013–5022 (2013).

33. Szabó, G. et al. Convergent patterns in the evolution of mealybug symbioses involving different intrabacterial symbionts. ISME J. 11, 715–726 (2017).

34. Takiya, D. M., Tran, P. L., Dietrich, C. H. & Moran, N. A. Co-cladogenesis spanning three phyla: Leafhoppers (*Insecta: Hemiptera: Cicadellidae*) and their dual bacterial symbionts. Mo.l Ecol. 15, 4175–4191 (2006).

35. Asnicar, F., et al. Studying Vertical Microbiome Transmission from Mothers to Infants by Strain-Level Metagenomic Profiling. mSystems 2, (2017).

36. Bobay, L.-M., Wissel, E. F. & Raymann, K. Strain Structure and Dynamics Revealed by Targeted Deep Sequencing of the Honey Bee Gut Microbiome. mSphere 5, (2020).

37. Sato, Y. et al. Fidelity varies in the symbiosis between a gutless marine worm and its microbial consortium. Microbiome 10, (2022).

38. Ferretti, P. et al. Mother-to-Infant Microbial Transmission from Different Body Sites Shapes the Developing Infant Gut Microbiome. Cell Host Microbe 24, 133–145.e5 (2018).

39. Rock, D. I. et al. Context-dependent vertical transmission shapes strong endosymbiont community structure in the pea aphid, *Acyrthosiphon pisum*. Mol. Eco.l 27, 2039–2056 (2018).

40. Busch, K. et al. Biodiversity, environmental drivers, and sustainability of the global deep-sea sponge microbiome. Nat. Commun. 13, (2022).

41. Quigley, K. M., Warner, P. A., Bay, L. K. & Willis, B. L. Unexpected mixed-mode transmission and moderate genetic regulation of *Symbiodinium* communities in a brooding coral. Heredity 121, 524–536 (2018).

42. Brooks, A. W., Kohl, K. D., Brucker, R. M., van Opstal, E. J. & Bordenstein, S. R. Phylosymbiosis: Relationships and Functional Effects of Microbial Communities across Host Evolutionary History. PLoS Biol. 14, (2016).

43. Brooks, A. W., Kohl, K. D., Brucker, R. M., van Opstal, E. J. & Bordenstein, S. R. Correction: Phylosymbiosis: Relationships and Functional Effects of Microbial Communities across Host Evolutionary History. PLoS Biol. 15, 1–2 (2017).

44. Pollock, F. J. et al. Coral-associated bacteria demonstrate phylosymbiosis and cophylogeny. Nat. Commun. 9, (2018).

45. Guo, W. P. et al. Extensive genetic diversity of *Rickettsiales* bacteria in multiple mosquito species. Sci. Rep. 6, (2016).

46. Cross, K. L. et al. Genomes of Gut Bacteria from *Nasonia* Wasps Shed Light on Phylosymbiosis and Microbe-Assisted Hybrid Breakdown. mSystems 6, (2021).

47. Gaulke, C. A. et al. Ecophylogenetics clarifies the evolutionary association between mammals and their gut microbiota. mBio 9, (2018).

48. O’Brien, P. A. et al. Diverse coral reef invertebrates exhibit patterns of phylosymbiosis. ISME J.14, 2211–2222 (2020).

49. Noda, S. et al. Cospeciation in the triplex symbiosis of termite gut protists (*Pseudotrichonympha* spp.), their hosts, and their bacterial endosymbionts. Mol. Ecol. 16, 1257–1266 (2007).

50. Qin, M., Chen, J., Xu, S., Jiang, L. & Qiao, G. Microbiota associated with *Mollitrichosiphum* aphids (*Hemiptera: Aphididae: Greenideinae*): diversity, host species specificity and phylosymbiosis. Environ. Microbiol. 23, 2184–2198 (2021).

51. Bourguignon, T. et al. Rampant Host Switching Shaped the Termite Gut Microbiome. Curr. Biol. 28, 649–654.e2 (2018).

52. Franzenburg, S., et al. Distinct antimicrobial peptide expression determines host species-specific bacterial associations. Proc. Natl. Acad. Sci. U.S.A 110, E3730 LP–E3738 (2013).

53. Tinker, K. A. & Ottesen, E. A. Phylosymbiosis across Deeply Diverging Lineages of Omnivorous Cockroaches (Order *Blattodea*). Appl. Environ. Microbiol. 86, e02513–19 (2020).

54. Sabrina Pankey, M., et al. Cophylogeny and convergence shape holobiont evolution in sponge– microbe symbioses. *Nat*. Ecol. Evol. 6, 750–762 (2022).

55. Dubilier, N. et al. Endosymbiotic sulphate-reducing and sulphide-oxidizing bacteria in an oligochaete worm. Nature 411, 298–302 (2001).

56. Bergin, C. et al. Acquisition of a Novel Sulfur-Oxidizing Symbiont in the Gutless Marine Worm *Inanidrilus exumae*. Appl. Environ. Microbiol. 84, e02267–17 (2018).

57. Dubilier, N. et al. Phylogenetic diversity of bacterial endosymbionts in the gutless marine oligochete *Olavius loisae* (Annelida). Mar. Ecol. Prog. Ser. 178, 271–280 (1999).

58. Dubilier, N., Giere, O., Distel, D. L. & Cavanaugh, C. M. Characterization of chemoautotrophic bacterial symbionts in a gutless marine worm (*Oligochaeta, Annelida*) by phylogenetic 16S rRNA sequence analysis and in situ hybridization. Appl. Environ. Microbiol. 61, 2346–2350 (1995).

59. Ruehland, C., Blazejak, A., Lott, C., Loy, A. & Erséus Christer & Dubilier, N. Multiple bacterial symbionts in two species of co-occurring gutless oligochaete worms from Mediterranean sea grass sediments. Environ. Microbiol. 10, 3404–3416 (2008).

60. Blazejak, A., Erseus, C. & Amann Rudolf & Dubilier, N. Coexistence of bacterial sulfide oxidizers, sulfate reducers, and spirochetes in a gutless worm (*Oligochaeta*) from the Peru margin. Appl. Environ. Microbiol. 71, 1553–1561 (2005).

61. Blazejak, A., Kuever, J., Erséus, C., Amann, R. & Dubilier, N. Phylogeny of 16S rRNA, Ribulose 1,5-Bisphosphate Carboxylase/Oxygenase, and Adenosine 5′-Phosphosulfate Reductase Genes from Gamma- and Alphaproteobacterial Symbionts in Gutless Marine Worms (Oligochaeta) from Bermuda and the Bahamas. Appl. Environ. Microbiol. 72, 5527–5536 (2006).

62. Bernot, J., et al. World Register of Marine Species (WoRMS). https://www.marinespecies.org (2025).

63. Woyke, T. et al. Symbiosis insights through metagenomic analysis of a microbial consortium. Nature 443, 950–955 (2006).

64. Erséus, C. et al. Phylogenomic analyses reveal a Palaeozoic radiation and support a freshwater origin for clitellate annelids. Zool. Scr. 49, 614–640 (2020).

65. Kleiner, M. et al. Use of carbon monoxide and hydrogen by a bacteria-animal symbiosis from seagrass sediments. Environ. Microbiol. 17, 5023–5035 (2015).

66. Kleiner, M. et al. Metaproteomics of a gutless marine worm and its symbiotic microbial community reveal unusual pathways for carbon and energy use. Proc. Natl. Acad. Sci. U.S.A 109, E1173 LP–E1182 (2012).

67. Zimmermann, J. et al. Closely coupled evolutionary history of ecto- and endosymbionts from two distantly related animal phyla. Mol. Eco.l 25, 3203–3223 (2016).

68. Giere, O. Ecology and Biology of Marine Oligochaeta – an Inventory Rather than another Review. Hydrobiologia 564, 103–116 (2006).

69. Giere, O. & Langheld, C. Structural organisation, transfer and biological fate of endosymbiotic bacteria in gutless oligochaetes. Mar. Biol. 93, 641–650 (1987).

70. Schimak, M. P. Transmission of Bacterial Symbionts in the Gutless Oligochaete Olavius algarvensis. PhD thesis, University of Bremen, Bremen (2015).

71. Krieger, J. Funktion und Übertragung endosymbiontischer Bakterien bei darmlosen marinen Oligochaeten. PhD thesis, University of Hamburg, Hamburg (2000).

72. Bergin, C. et al. Acquisition of a Novel Sulfur-Oxidizing Symbiont in the Gutless Marine Worm *Inanidrilus exumae*. Appl. Environ. Microbiol. 84, e02267–17 (2018).

73. Dubilier, N. & Blazejak Anna & Rühland C. Symbioses between Bacteria and Gutless Marine Oligochaetes. Molecular Basis of Symbiosis 251–275 (Springer, Berlin, Heidelberg, 2006).

74. Seah, B. K. B. et al. Sulfur-Oxidizing Symbionts without Canonical Genes for Autotrophic CO2 Fixation. mBio 10, e01112–19 (2019).

75. Dubilier, N. & Bergin Claudia & Lott, C. Symbiotic diversity in marine animals: the art of harnessing chemosynthesis. Nat. Rev. Micro. 6, 725–740 (2008).

76. Nakabachi, A. et al. Transcriptome analysis of the aphid bacteriocyte, the symbiotic host cell that harbors an endocellular mutualistic bacterium, Buchnera. Proc. Natl. Acad. Sci. U.S.A. 102, 5477 LP –5482 (2005).

77. Login, F. H. et al. Antimicrobial Peptides Keep Insect Endosymbionts Under Control. Science 334, 362 LP –365 (2011).

78. McFall-Ngai, M. Care for the community. Nature 445, 153 (2007).

79. Anselme, C. et al. Identification of the Weevil immune genes and their expression in the bacteriome tissue. BMC Biol. 6, 43 (2008).

80. Mazel, F. et al. Is Host Filtering the Main Driver of Phylosymbiosis across the Tree of Life? mSystems 3, e00097–18 (2018).

81. Mallott, E. K. & Amato, K. R. Host specificity of the gut microbiome. Nat. Rev. Microbiol. 19, 639–653 (2021).

82. Bowers, R. M. et al. Minimum information about a single amplified genome (MISAG) and a metagenome-assembled genome (MIMAG) of bacteria and archaea. Nat. Biotechnol. 35, 725–731 (2017).

83. Jain, C., Rodriguez-R, L. M., Phillippy, A. M., Konstantinidis, K. T. & Aluru, S. High throughput ANI analysis of 90K prokaryotic genomes reveals clear species boundaries. Nat. Commun. 9, (2018).

84. Bright, M. & Bulgheresi, S. A complex journey: transmission of microbial symbionts. Nat. Rev. Microbiol. 8, 218–230 (2010).

85. Russell, S. L. Transmission mode is associated with environment type and taxa across bacteria-eukaryote symbioses: a systematic review and meta-analysis. FEMS Microbiol. Lett. 366, (2019).

86. Matumba, T. G., Oliver, J., Barker, N. P., McQuaid, C. D. & Teske, P. R. Intraspecific mitochondrial gene variation can be as low as that of nuclear rRNA. F1000Res 9, (2020).

87. Xu, J. The inheritance of organelle genes and genomes: patterns and mechanisms. Genome 48, 951–958 (2005).

88. Ishikawa, S. A., Zhukova, A., Iwasaki, W. & Gascuel, O. A Fast Likelihood Method to Reconstruct and Visualize Ancestral Scenarios. Mol. Biol. Evol. 36, 2069–2085 (2019).

89. Mazel, F., Prasad, A. & Engel, P. Host specificity of gut microbiota associated with social bees: patterns and processes. Microbiol. Mol Biol. Rev. 89 (2025)

90. Moeller, A. H. Partner fidelity, not geography, drives co-diversification of gut microbiota with hominids. Biol. Lett. 21, 20240454 (2025).

91. Bongrand, C. et al. Evidence of Genomic Diversification in a Natural Symbiotic Population Within Its Host. Front. Microbiol. 13, (2022).

92. Sachs, J. L. & Simms, E. L. Pathways to mutualism breakdown. Trends Ecol. Evol. 21, 585–592 (2006).

93. Toby Kiers, E., Palmer, T. M., Ives, A. R., Bruno, J. F. & Bronstein, J. L. Mutualisms in a changing world: an evolutionary perspective. Ecol. Lett. 13, 1459–1474 (2010).

94. Russell, S. L. et al. Horizontal transmission and recombination maintain forever young bacterial symbiont genomes. PLoS Genet. 16, e1008935 (2020).

95. Kleiner, M. et al. Metaproteomics method to determine carbon sources and assimilation pathways of species in microbial communities. Proc. Natl. Acad. Sci. U.S.A. 115, E5576 LP–E5584 (2018).

96. Rühlemann, M. C. et al. Functional host-specific adaptation of the intestinal microbiome in hominids. Nat. Commun. 15, 326 (2024).

97. Kwong, W. K. et al. Dynamic microbiome evolution in social bees. Sci. Adv. 3, e1600513 (2025).

98. Quast, C. et al. The SILVA ribosomal RNA gene database project: improved data processing and web-based tools. Nucleic Acids Res. 41, D590–6 (2013).

99. Yilmaz, P. et al. The SILVA and “All-species Living Tree Project (LTP)” taxonomic frameworks. Nucleic Acids Res. 42, D643–D648 (2014).

100. Bankevich, A. et al. SPAdes: a new genome assembly algorithm and its applications to single-cell sequencing. J. Comput. Biol. 19, 455–477 (2012).

101. Gruber-Vodicka, H. R., Seah, B. K. B. & Pruesse, E. phyloFlash: Rapid Small-Subunit rRNA Profiling and Targeted Assembly from Metagenomes. mSystems 5, e00920–20 (2020).

102. Katoh, K., Kuma, K., Toh, H. & Miyata, T. MAFFT version 5: improvement in accuracy of multiple sequence alignment. Nucleic Acids Res. 33, 511–518 (2005).

103. Katoh, K. & Standley, D. M. MAFFT Multiple Sequence Alignment Software Version 7: Improvements in Performance and Usability. Mol. Biol. Evol. 30, 772–780 (2013).

104. Katoh, K., Misawa, K., Kuma, K. & Miyata, T. MAFFT: a novel method for rapid multiple sequence alignment based on fast Fourier transform. Nucleic Acids Res. 30, 3059–3066 (2002).

105. Letunic, I. & Bork, P. Interactive Tree Of Life (iTOL): an online tool for phylogenetic tree display and annotation. Bioinformatics 23, 127–128 (2007).

106. Paradis, E., Claude, J. & Strimmer, K. APE: Analyses of Phylogenetics and Evolution in R language. Bioinformatics 20, 289–290 (2004).

107. Edgar, R. C. Search and clustering orders of magnitude faster than BLAST. Bioinformatics 26, 2460–2461 (2010).

108. Pruesse, E. & Peplies Jörg & Glöckner, F. O. SINA: Accurate high-throughput multiple sequence alignment of ribosomal RNA genes. Bioinformatics 28, 1823–1829 (2012).

109. Price, M. N., Dehal, P. S. & Arkin, A. P. FastTree 2 – Approximately Maximum-Likelihood Trees for Large Alignments. PLoS One 5, e9490 (2010).

110. Miller, C. S., Baker, B. J., Thomas, B. C. & Singer Steven W. & Banfield, J. F. EMIRGE: reconstruction of full-length ribosomal genes from microbial community short read sequencing data. Genome Biol. 12, R44 (2011).

111. Pruitt, K. D. & Maglott, D. R. RefSeq and LocusLink: NCBI gene-centered resources. Nucleic Acids Res. 29, 137–140 (2001).

112. Altschul, S. F., Gish, W., Miller, W. & Myers E. W. & Lipman, D. J. Basic local alignment search tool. J. Mol. Biol. 215, 403–410 (1990).

113. Bartram, A. K. et al. Exploring links between pH and bacterial community composition in soils from the Craibstone Experimental Farm. FEMS Microbiol. Ecol. 87, 403–415 (2014).

114. Wickham, H. The Split-Apply-Combine Strategy for Data Analysis. J. Stat. Softw. 40, (2011).

115. Venables, W. N. & Ripley, B. D. Modern Applied Statistics with S. (Springer, 2002).

116. Wickham, H. et al. Welcome to the Tidyverse. J. Open Source Softw. 4, 1686 (2019).

117. Mantel, N. The Detection of Disease Clustering and a Generalized Regression Approach. Cancer Res. 27, 209 LP –220 (1967).

118. Anderson, M. J. A new method for non-parametric multivariate analysis of variance. Austral. Ecol. 26, 32–46 (2001).

119. Csardi, G. & Nepusz, T. The igraph software package for complex network research. *InterJournal*, Complex Syst.1695, 1–9 (2006).

120. Robinson, D. F. & Foulds, L. R. Comparison of phylogenetic trees. Math. Biosci. 53, 131–147 (1981).

121. Bogdanowicz, D., Giaro, K. & Wróbel, B. TreeCmp: Comparison of Trees in Polynomial Time. Evol. Bioinfor. 8, 475–487 (2012).

122. Bogdanowicz, D. & Giaro, K. On a matching distance between rooted phylogenetic trees. Int. J. Appl. Math. Comput. Sci. 23, 669–684.

122. Chklovski, A., Parks, D. H., Woodcroft, B. J. & Tyson, G. W. CheckM2: a rapid, scalable and accurate tool for assessing microbial genome quality using machine learning. Nat. Methods. 20, 1203–1212 (2023).

123. Laslett, D. & Canback, B. ARAGORN, a program to detect tRNA genes and tmRNA genes in nucleotide sequences. Nucleic Acids Res. 32, 11–16 (2004).

124. Parks, D. H. et al. A complete domain-to-species taxonomy for Bacteria and Archaea. Nat. Biotechnol. 38, 1079–1086 (2020).

125. Parks, D. H. et al. A standardized bacterial taxonomy based on genome phylogeny substantially revises the tree of life. Nat. Biotechnol. 36, 996–1004 (2018).

126. Chaumeil, P.-A., Mussig, A. J., Hugenholtz, P. & Parks, D. H. GTDB-Tk: a toolkit to classify genomes with the Genome Taxonomy Database. Bioinformatics 36, 1925–1927 (2019).

127. Verdes, A., et al. Molecular phylogeny of *Odontosyllis* (*Annelida, Syllidae*): A recent and rapid radiation of marine bioluminescent worms. bioRxiv 241570 (2018). doi:10.1101/241570.

128. Anderson, F. E. et al. Phylogenomic analyses of *Crassiclitellata* support major Northern and Southern Hemisphere clades and a Pangaean origin for earthworms. BMC Evol. Biol. 17, 123 (2017).

129. Bouckaert, R. et al. BEAST 2: A Software Platform for Bayesian Evolutionary Analysis. PLoS Comput. Biol. 10, e1003537 (2014).

130. Hugoson, E., Ammunét, T. & Guy, L. Host-adaptation in Legionellales; is 2.4 Ga, coincident with eukaryogenesis. bioRxiv 852004 (2020). doi:10.1101/852004

