## Supplementary Material for "Repeated losses and gains of bacterial symbionts in gutless marine annelids over 150 million years"

### Contents

Contains or links to
Supplementary Tables 1-11; provided in a separate file
Supplementary Figures 1-43; Figures S2-34 are shared as links to digital phylogenies
Supplementary Notes 1-5

**Tables**

S1: Sample overview and contextual data

S2: Taxonomy of symbiont clades

S3: Relative symbiont abundances in gutless oligochaete samples

S4: Mean abundance and standard deviation of each symbiont genus in each host species.

S5: Estimated genome statistics of symbiont MAGs.

S6 Average nucleotide identity (ANI) of symbiont individuals from the same genus.

S7: List of symbiont species and their host range. Symbiont species highlighted in bold were shared by two or more host species. Yellow highlights cases where individuals from the same host species were associated with different symbiont species from the same genus.

S8: Examples of host specimen from different sites that harbor different symbiont species. Cases where specimens from different locations were associated with the same symbiont species are highlighted in bold.

S9: Numbers of shared nodes between the host 28S rRNA and the mtCOI phylogenies.

S10: Estimated acquisitions and losses of symbiont clades based on host nuclear (28 rRNA) lineages

S11: Library preparation, read length and number of reads

#### Figures

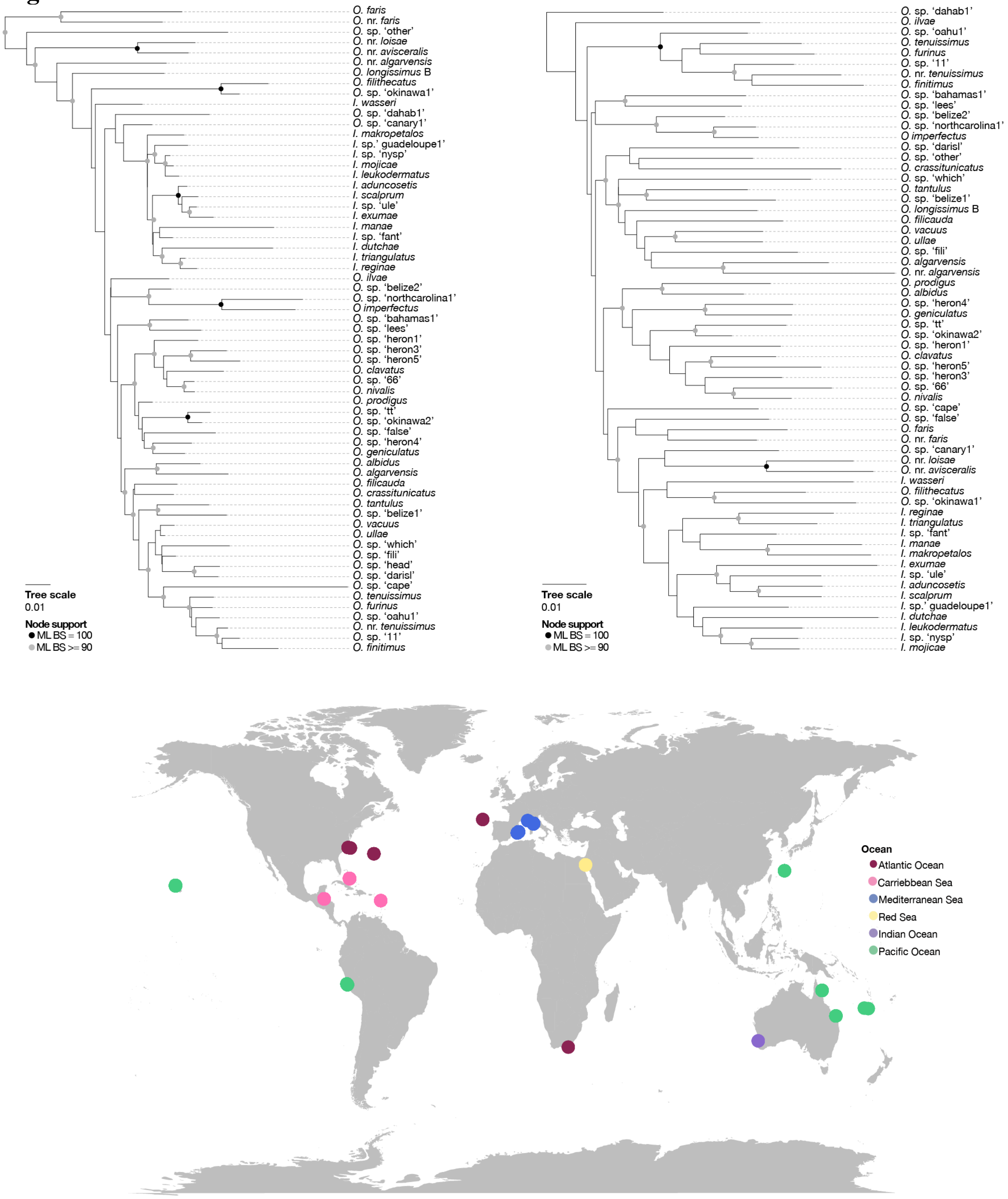


**Figure S1: Host 28S rRNA phylogeny (A), host mtCOI phylogeny (B) and sampling locations (C).** Maximum-likelihood phylogeny of both genes based on MAFFT alignments and calculated with IQ-TREE using automated model selection. Node support is based on 1000 bootstrap iterations.

**Figure S2-S34: 16S rRNA phylogenies of the symbionts of gutless oligochaetes from each genus and their relatives from public databases.** Sequences obtained in this study are highlighted in bold, black letters. Sequences are color-coded as follows: black, bold: symbionts of gutless oligochaetes, this study; red, bold: symbionts of gutless oligochaetes, previous studies; pink: symbionts from other chemosymbiotic hosts (stilbonematine nematodes, *Astomonema* nematodes, *Kentrophoros* ciliates, lucinid clams); orange: coral-associated bacteria; purple: sponge-associated bacteria; blue: bacteria associated with other marine animals (dolphins, fish gut). All sequences from environmental or unknown sources are shown in gray. Type strains are highlighted in bold gray letters.

**Figure S2: 16S phylogeny of Actionmarinales1 symbionts and their relatives.**
https://itol.embl.de/tree/10111897188401745915130

**Figure S3: 16S phylogeny of Alpha1 symbionts and their relatives.**
https://itol.embl.de/tree/10111897191271745915136

**Figure S4: 16S phylogeny of Alpha2 symbionts and their relatives.**
https://itol.embl.de/tree/10111897195631745915142

**Figure S5: 16S phylogeny of Alpha3 symbionts and their relatives.**
https://itol.embl.de/tree/10111897211711745915155

**Figure S6: 16S phylogeny of Alpha4 symbionts and their relatives.**
https://itol.embl.de/tree/10111897228231745915171

**Figure S7: 16S phylogeny of Alpha5 symbionts and their relatives.**
https://itol.embl.de/tree/10111897235721745915178

**Figure S8: 16S phylogeny of Alpha6 symbionts and their relatives.**
https://itol.embl.de/tree/10111897243951745915184

**Figure S9: 16S phylogeny of Alpha7 symbionts and their relatives.**
https://itol.embl.de/tree/10111897268891745915203

**Figure S10: 16S phylogeny of Alpha8 symbionts and their relatives.**
https://itol.embl.de/tree/10111897268891745915203

**Figure S11: 16S phylogeny of Alpha9 symbionts and their relatives.**
https://itol.embl.de/tree/10111897274501745915210

**Figure S12: 16S phylogeny of Alpha10 symbionts and their relatives.**
https://itol.embl.de/tree/10111897279131745915216

**Figure S13: 16S phylogeny of Alpha12 symbionts and their relatives.**
https://itol.embl.de/tree/10111897282671745915222

**Figure S14: 16S phylogeny of Alpha13 symbionts and their relatives.**
https://itol.embl.de/tree/10111897291051745915230

**Figure S15: 16S phylogeny of Alpha14 symbionts and their relatives.**
https://itol.embl.de/tree/10111897299021745915236

**Figure S16: 16S phylogeny of Alpha15 symbionts and their relatives.**
https://itol.embl.de/tree/10111897312261745915247

**Figure S17: 16S phylogeny of Alpha16 symbionts and their relatives.**
https://itol.embl.de/tree/10111897317581745915254

**Figure S18: 16S phylogeny of Delta1 symbionts and their relatives.**
https://itol.embl.de/tree/10111897323961745915260

**Figure S19: 16S phylogeny of Delta2 symbionts and their relatives.**
https://itol.embl.de/tree/10111897332101745915266

**Figure 20: 16S phylogeny of Delta3 symbionts and their relatives.**
https://itol.embl.de/tree/10111897334451745915272

**Figure S21: 16S phylogeny of Delta4 symbionts and their relatives.**
https://itol.embl.de/tree/10111897340421745915277

**Figure S22: 16S phylogeny of Delta5 symbionts and their relatives.**
https://itol.embl.de/tree/10111897346271745915283

**Figure S23: 16S phylogeny of Delta11 symbionts and their relatives.**
https://itol.embl.de/tree/10111897354021745915290

**Figure S24: 16S phylogeny of Delta12 symbionts and their relatives.**
https://itol.embl.de/tree/10111897361711745915297

**Figure S25: 16S phylogeny of Delta13 symbionts and their relatives.**
https://itol.embl.de/tree/10111897367611745915303

**Figure S26: 16S phylogeny of Delta14 symbionts and their relatives.**
https://itol.embl.de/tree/10111897379891745915319

**Figure S27: 16S phylogeny of *Thiosymbion* (Gamma1) symbionts and their relatives.**
https://itol.embl.de/tree/10111897388531745915325

**Figure S28: 16S phylogeny of Gamma2 symbionts and their relatives.**
https://itol.embl.de/tree/10111897395921745915333

**Figure S29: 16S phylogeny of Gamma3 symbionts and their relatives.**
https://itol.embl.de/tree/10111897407511745915344

**Figure S30: 16S phylogeny of Gamma4 symbionts and their relatives.**
https://itol.embl.de/tree/10111897413931745915350

**Figure S31: 16S phylogeny of Gamma5 symbionts and their relatives.**
https://itol.embl.de/tree/10111897427471745915364

**Figure S32: 16S phylogeny of Gamma7 symbionts and their relatives.**
https://itol.embl.de/tree/10111897434051745915369

**Figure S33: 16S phylogeny of Marinimicrobia1 symbionts and their relatives.**
https://itol.embl.de/tree/10111897444071745915377

**Figure S34: 16S phylogeny of Spiro1 symbionts and their relatives.**
[https://itol.embl.de/tree/10111897453271745915385](https://itol.embl.de/tree/10111897453271745915385)

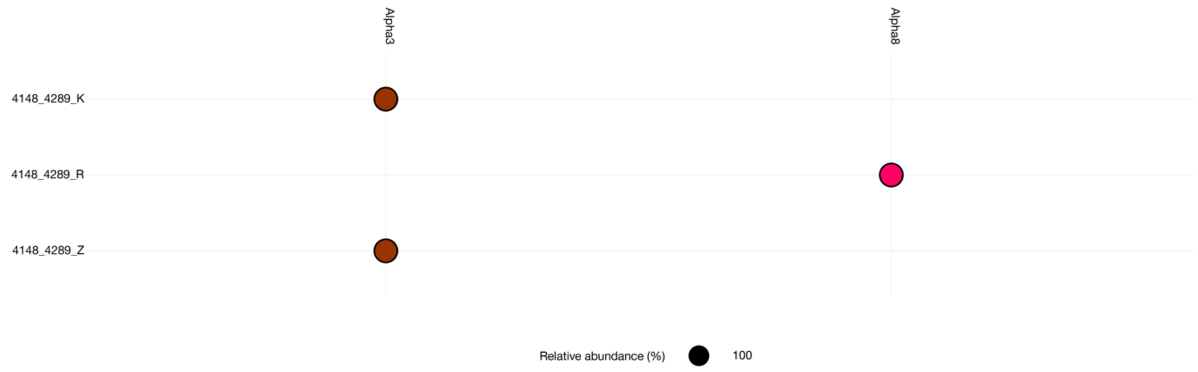


**Figure S35: Abundance plot for bacterial genera associated with gut-bearing relatives:** Relative abundances estimated with EMIRGE.

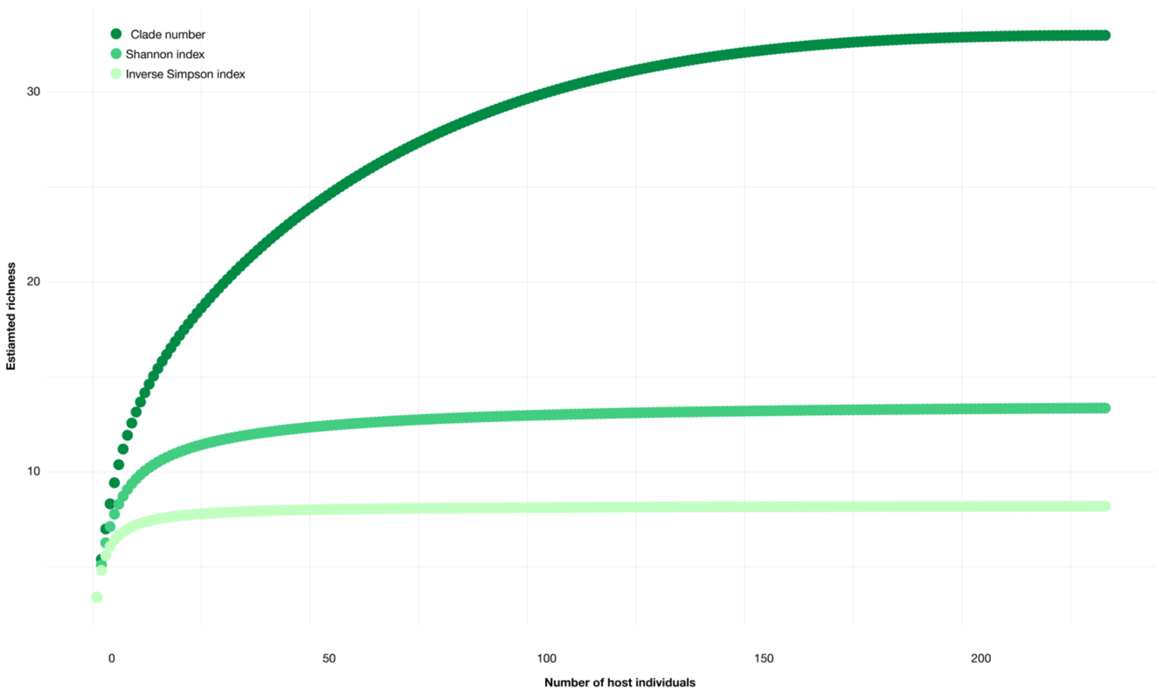


**Figure S36: Gutless oligochaete symbionts belong to a limited pool of bacterial genera.** Rarefaction analysis of the number of symbiont genera over the number of analyzed host individuals. Richness of symbiont genera (‘richness’) was analyzed as absolute number of genera as well as the Shannon Index and the Inverse Simson Index.

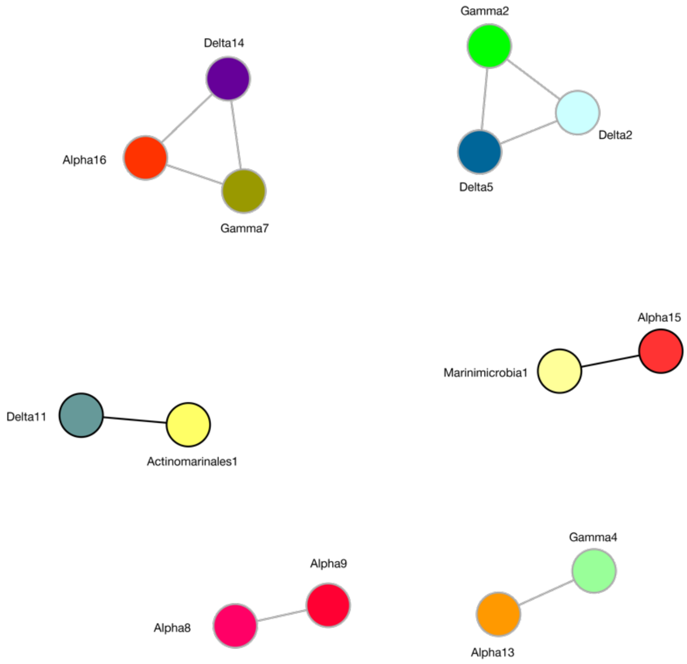


**Figure S37: Network showing significant co-occurrences of symbiont genera.** Unbiased correlation network analysis based on relative abundance of symbiont genera per host species as estimated with EMIRGE. Co-occurrences were calculated using Spearman's correlations and corrected using the Benjamini-Hochberg standard false discovery rate correction. Networks of significant co-occurrences were generated using igraph[1]**.** Co-occurrences highlighted by black lines were found across two host species, co-occurrences highlight with gray lines were found within a single host species.

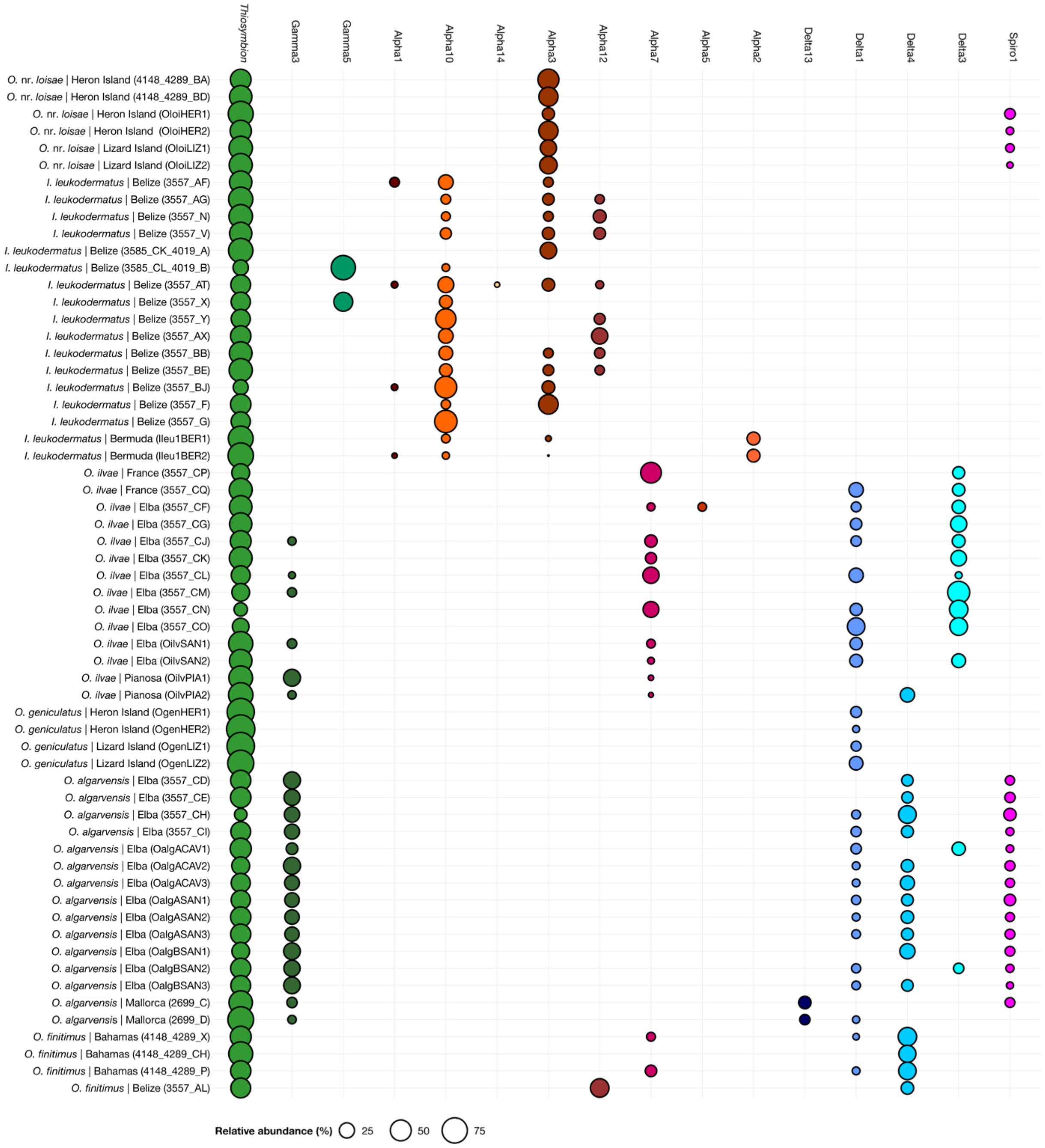


**Figure S38: Relative abundances of symbiont genera in specimens from the same host species, sampled in different locations.** Relative abundances were estimated with EMIRGE.

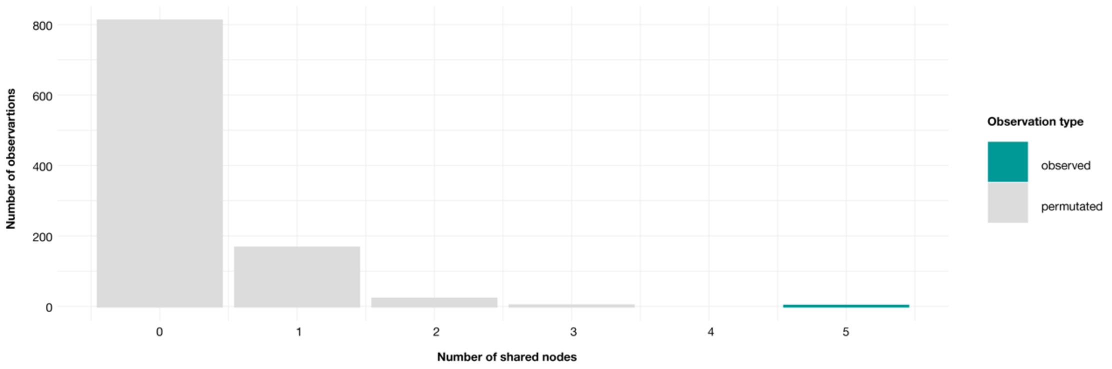


**Figure S39: Test for phylosymbiosis.** Number of shared nodes between the dendrogram reflecting symbiont community composition and the host phylogeny (green) in comparison to the number of shared nodes between 1000 dendrograms derived from permutated versions of the symbiont abundance data and the host phylogeny (gray).

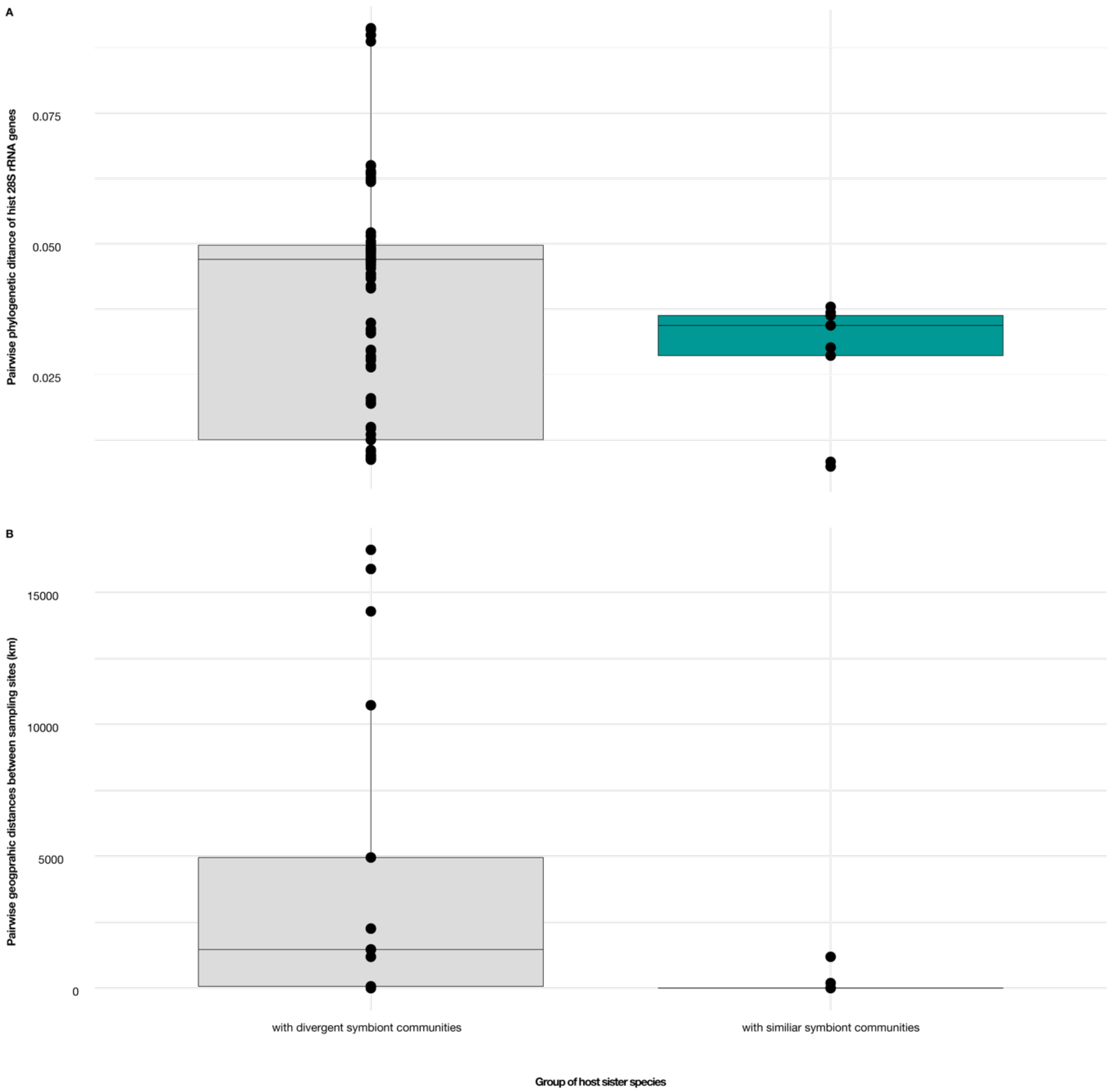


**Figure S40: Host sister species that share similar symbiont communities are more closely related and occur in closer geographic proximity.** A: pairwise phylogenetic distance between host individuals from sister species based on their 28S rRNA phylogeny. B: pairwise geographic distance between host individuals from sister species based on their sampling location.

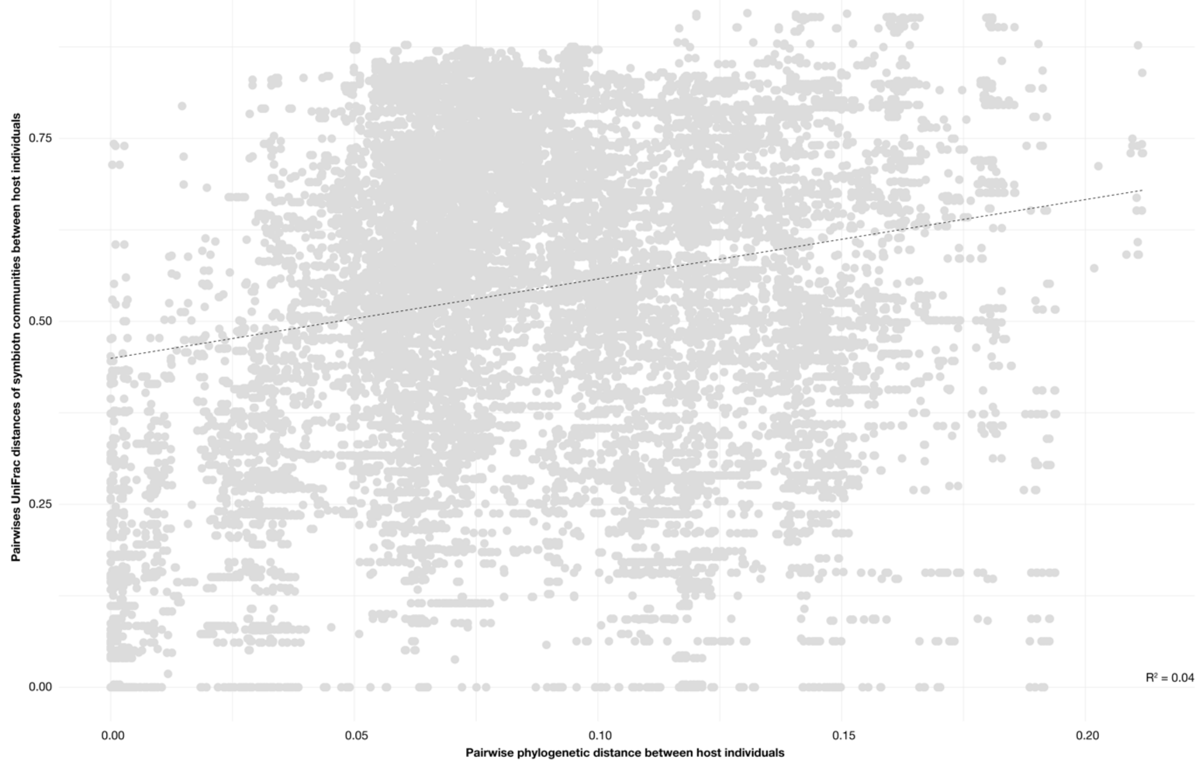


**Figure S41: Host phylogeny does not explain symbiont community composition.** Pairwise UniFrac dissimilarities of symbiont community composition in relation to the pairwise nucleotide dissimilarity of the host 28S rRNA genes.

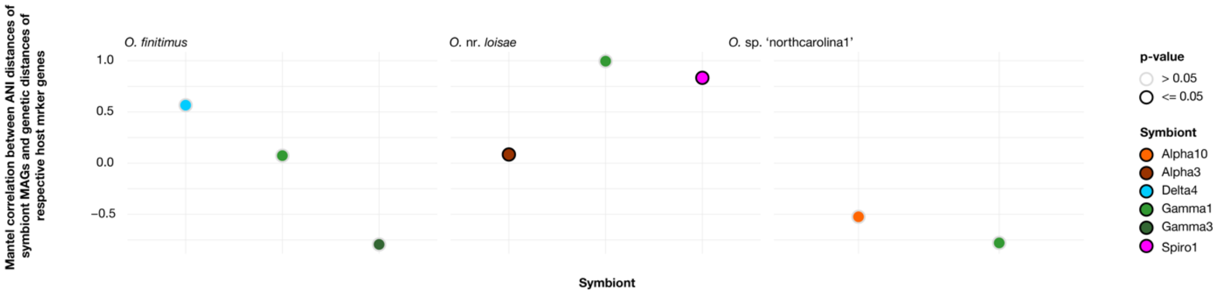


**Figure S42: Partner fidelity in host species where the analyses of symbiont fidelity could only be performed for the 28S rRNA gene.** Each panel shows the correlation values for the indicated host species between symbiont ANIs and the host nuclear gene 28S rRNA. For all data points in this image, MAGs for each symbiont genus were recovered from at least three individuals per host species. Significant correlations (Mantel test; p ≤0.05) are highlighted.

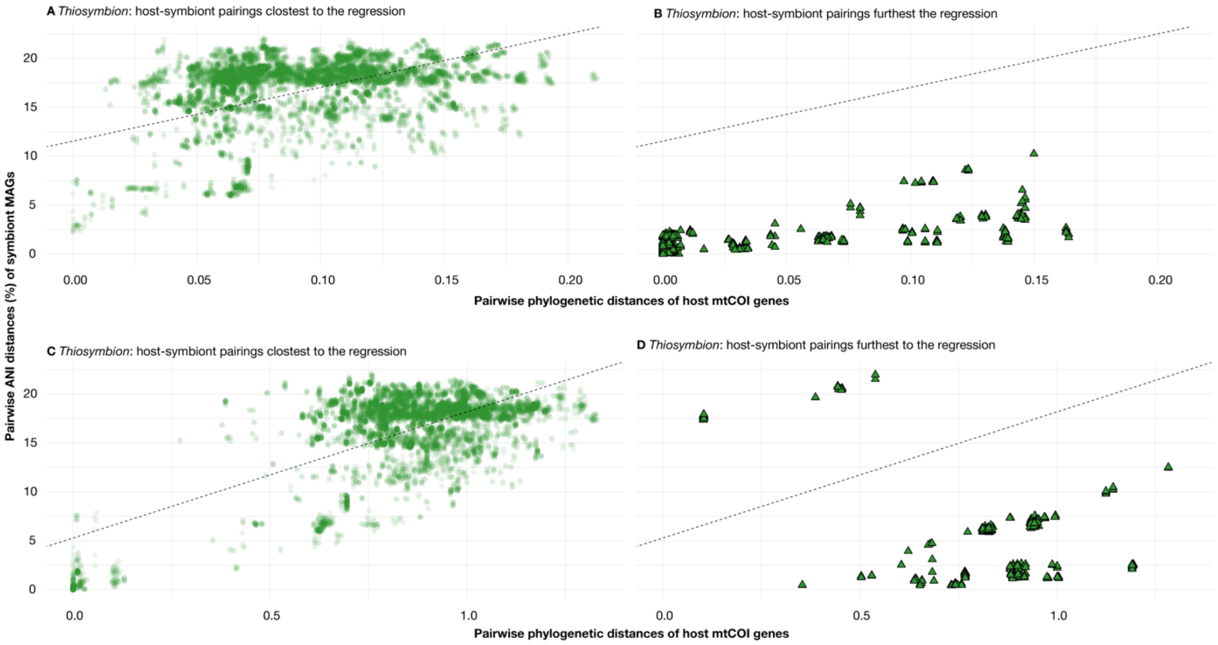


**Figure S43: *Thiosymbion* – host codivergence for the 28S rRNA and mtCOI gene.** Pairwise ANI dissimilarity from MAGs of *Thiosymbion* with the corresponding pairwise genetic dissimilarity of host marker genes, split by perpendicular distance from the regression.

#### Notes

**Note S1: Curation of symbiont clades**

To minimize the risk of contamination in our dataset, we retained only those clades in downstream analyses that were present in at least 50% of individuals from a given host species and sampling site, with a minimum of two host individuals per host species and site. For host species that were sampled at two locations, we retained symbiont clades using the same criteria and were found in one of the two sampling locations. Clades found exclusively in libraries produced at one sequencing center—without consistent presence in the same host species sequenced at another sequencing center—were also excluded.

**Note S2: Phylogenetic relationships of gutless oligochaete symbiont clades**

To identify other hosts or environments that symbiont genera were associated with, we screened publicly available 16S rRNA gene sequence data, and analyzed the phylogenetic relations between the symbionts of gutless oligochaetes and their closest relatives (Fig. S4-S36). We found that 21 of the 33 symbiont genera consisted exclusively of gutless oligochaete symbionts. Two of the remaining 12 clades represented sister clades to phylotypes also detected in Stilbonematinae nematodes (Gamma5 and Gamma7). In addition, two symbiont clades were phylogenetically intermixed with characterized symbionts from animals of co-occurring meiofauna: as previously reported, *Thiosymbion* was also associated with Stilbonematinae and *Astomonema* nematodes. Additionally, the Gamma4 symbionts were already described as '*Ca.* Kentron' symbionts of *Kentrophoros* ciliates. The symbionts from the remaining six clades were phylogenetically intermixed with bacteria that were previously detected in environmental samples (Alpah14, Delta1, Delta3, Delta4, and Gamma3). The degree to which these gutless oligochaete symbionts were phylogenetically intermixed with free-living relatives varied between clades and could point towards different modes of acquisition and inheritance.

To understand whether gutless oligochaete symbiont clades were associated specifically with gutless hosts or also occurred in related marine oligochaetes, we screened nine specimens that were morphologically identified as members of the closely related gut-bearing *Phallodrilinae*. Two symbiont clades, the Alpha3 and Alpha8, could also be detected in these gut-bearing relatives, indicating that most of the symbiont clades are specific to gutless oligochaetes (Figure S36). Based on these results, we grouped the symbiont clades into three categories: i) symbionts only associated with gutless oligochaetes, ii) symbionts also associated with other marine invertebrates and iii) symbionts that were phylogenetically intermixed with populations of free-living bacteria.

**Note S3: Detection of a single 16S rRNA sequence of a free-living bacterium in the Spiro1 symbiont clade and a single 16S rRNA sequences from a coral in the *Thiosymbion* clade**

In the main text, we reported Spiro1 as a symbiont clade composed exclusively of sequences from gutless oligochaetes. However, during phylogenetic tree reconstruction, we identified one sequence (AQWX01000128) from a free-living bacterium that fell within the Spiro1 clade. Upon further investigation, we found that this sequence had been annotated as a *Staphylococcus* species by the researchers that had submitted the sequence to the public databases, without any additional metadata. Due to phylogenetic inconsistencies and the lack of information about the source of this sequence, we excluded it from further consideration.

Similarly, we identified a 16S rRNA gene sequence (GU118064), obtained from a coral sample within the *Thiosymbion* clade. Given that *Thiosymbion* hosts including gutless oligochaetes are commonly found in coral reef habitats and that *Thiosymbion* sequences were not detected in any of the many other coral samples analyzed, we infer that this sequence likely originated from a *Thiosymbion* host that was sampled with the coral. Based on its phylogenetic position, the host likely was a Stilbonematinae nematode of the genus *Eubostrichus*.

**Note S4: Symbiont-symbiont interactions**

In addition to host traits, symbiont-symbiont interactions may influence community composition as demonstrated in other highly specific symbiont consortia, such as those associated with plant hosts[2]. To investigate this possibility, we conducted an unbiased network analysis to test for patterns of co-occurrence and mutual exclusion among symbiont clades. Significant co-occurrences were calculated using Spearman's correlations and corrected using the Benjamini-Hochberg standard false discovery rate correction. Networks of significant co-occurrences were generated using igraph[1]. We found no evidence of symbiont exclusion, suggesting that community composition is not shaped by symbiont–symbiont competition. However, we identified six instances of co-occurrence: four involved symbiont clades co-existing within a single host species, while the remaining two were observed in clades co-occurring in closely related host sister species. The limited number of stable positive associations, confined to symbionts within a single host lineage, suggests that symbiont–symbiont interactions play only a minor role in structuring the symbiotic communities of gutless oligochaetes.

**Note S5: Distribution of symbiont species in across host species**

Overall, we observed specific associations between individuals of the same host species and symbiont species of the different genera, i.e., usually all individuals of one host species were associated with the same symbiont species. Host individuals of different species were associated with different symbiont species. This observation was true for 116 out of 136 analyzed symbiont species (Table S7). The remaining 20 symbiont species were shared by individuals of different species. These individuals were either from closely related host species or occurred in the same geographic region (Fig. S1, Tables S1, S3).

Similarly, we mostly observed that all individuals from one host species were associated with the same symbiont species. Exceptions were individuals of *O. clavatus* that were associated with two species of the Actinomarinales1 symbiont; individuals of *O.* nr. *loisae* that were associated with two species of the Alpha3 symbiont; individuals of *O. finitimus* that were associated with two species of the Alpha7 symbiont; individuals of *I. triangulatus* that were associated with two species of the Alpha10 symbiont; individuals of *I. dutchae* that were associated with four species of the Delta4 symbiont; individuals of *O.* sp. 'fili' that were associated with two species of *Thiosymbion*; and individuals of *O. algarvensis* that were associated with two species of *Thiosymbion* and the Gamma3 symbiont, respectively. Some of these exceptions are linked to the individuals of one species occurring at different sampling location (Table S7). These cases are further discussed in the main text and Table S8.

#### Extended data

All phylogenetic trees that were generated in this study are deposited here:
Symbiont 16S rRNA tree: https://itol.embl.de/tree/10111897149301745914966
Host 28S rRNA tree: https://itol.embl.de/tree/10111897153601745914987
Host mtCOI tree: https://itol.embl.de/tree/10111897152591745914979

### Literature

1. Csardi, G. & Nepusz, T. The igraph software package for complex network research. *InterJournal, Complex Sys*t. **1695**, 1–9 (2006).
2. Uroz, S., Courty, P. E. & Oger, P. Plant Symbionts Are Engineers of the Plant-Associated Microbiome. *Trends Plant. Sci.* **24**, 905–916 (2019).
